# Shoot and root thermomorphogenesis are linked by a developmental trade-off

**DOI:** 10.1101/2020.05.07.083246

**Authors:** Christophe Gaillochet, Yogev Burko, Matthieu Pierre Platre, Ling Zhang, Jan Simura, Vinod Kumar, Karin Ljung, Joanne Chory, Wolfgang Busch

## Abstract

Temperature is one of the most impactful environmental factors to which plants adjust their growth and development. While the regulation of temperature signaling has been extensively investigated for the aerial part of plants, much less is known and understood about how roots sense and modulate their growth in response to fluctuating temperatures. Here we found that shoot and root growth responses to high ambient temperature are coordinated during early seedling development. A shoot signaling module that includes HY5, the phytochromes and the PIFs exerts a central function in coupling these growth responses and control auxin levels in the root. In addition to the HY5/PIF-dependent shoot module, a regulatory axis composed of auxin biosynthesis and auxin perception factors controls root responses to high ambient temperature. Together, our findings show that shoot and root developmental responses to temperature are tightly coupled during thermomorphogenesis and suggest that roots integrate energy signals with local hormonal inputs.

## Introduction

Over the course of their life, plants are subjected to constant environmental fluctuations. Consequently, plants have evolved tremendous developmental plasticity that allows them to precisely adjust their development to environmental conditions and therefore to thrive in dynamically and often unpredictably changing environments. In particular, the early stage of seedling development constitutes a critical moment at which plants need to sense their environment and respond quickly to fine-tune their developmental programs and successfully establish themselves as autotrophic seedlings (reviewed in (Ha et al., 2017)). Not surprisingly, early life stages have been shown to strongly contribute to local adaptation (reviewed in (Donohue et al., 2010)).

Temperature is a pervasive environmental parameter influencing biological systems at all scales from the rate of biochemical reactions to the timing of developmental transitions (reviewed in (Penfield, 2008)). In addition, temperature shows important geographical, diurnal as well as seasonal variation. Importantly, plants are equipped with sophisticated molecular machineries to perceive temperature fluctuations, which allows them to sense and translate these signals into appropriate developmental responses. Accordingly, raising ambient temperature leads to increased elongation of the hypocotyl and root –a process called thermomorphogenesis (reviewed in (Quint et al., 2016)).

The molecular mechanisms underlying shoot thermo-responses have been largely investigated (Quint et al., 2016). In this context, the photoreceptor PHYTOCHROME B (PHYB) enables perception of higher ambient temperature by switching from an active to an inactive form (Legris et al., 2016). This process of phytochrome thermal reversion subsequently prevents sequestration and degradation of transcription factors such as the PHYTOCHROME INTERACTING FACTORs (PIFs) that can accumulate and promote the expression of downstream regulatory genes (Jung et al., 2016; Kumar et al., 2012; Park et al., 2018).

Among the PIF clade, PIF4 acts as a central signalling hub to mediate shoot thermomorphogenesis (Quint et al., 2016; Koini et al., 2009). Upon higher ambient temperature, PIF4 directly positively regulates the expression of a battery of genes including auxin biosynthetic genes *YUCCA8 (YUC8)* and *TRYPTOPHAN AMINOTRANSFERASE OF ARABIDOPSIS1* (*TAA1*), thereby promoting an elevation of auxin levels and increased hypocotyl cell elongation (Franklin et al., 2011; Sun et al., 2012). This regulatory circuit also integrates inputs from the transcription factor LONG HYPOCOTYL5 (HY5) that can act antagonistically to PIF4 by repressing *PIF4* expression or by directly regulating key PIF4 target genes including *YUC8* (Delker et al., 2014; Gangappa and Kumar, 2017). Both HY5 and PIF4 expression levels and protein abundance are tightly regulated by a plethora of factors (reviewed in (Lau and Deng, 2012; Quint et al., 2016)). Among those, CONSTITUTIVE PHOTOMORPHOGENESIS PROTEIN1 (COP1) and DEETIOLATED1 (DET1) trigger HY5 degradation and promote both *PIF4* expression and protein stabilization (Gangappa and Kumar, 2017; Osterlund et al., 2000; Saijo et al., 2003; Yanagawa et al., 2004). The collective genetic activity of *PIF4, HY5, COP1* and *DET1* defines an intertwined regulatory module that acts at the interface between light and temperature signalling (Delker et al., 2014; Gangappa and Kumar, 2017). Interestingly, HY5 protein has also been shown to translocate from the shoot to the root and to coordinate carbon fixation with nitrogen uptake (Chen et al., 2016).

Importantly, roots can autonomously sense and respond to temperature (Bellstaedt et al., 2019), which might allow them to reach deeper and cooler layers of the soil under warm surface conditions (Illston and Fiebrich, 2017). However, in contrast to the shoot, the molecular mechanisms underlying plant root thermo-responses have so far remained elusive. Similarly to the shoot, maintenance of auxin homeostasis is critical for the root response to temperature (Wang et al., 2016). In line with this idea, auxin signaling increases upon perception of higher ambient temperature (Hanzawa et al., 2013; Wang et al., 2016). In this context the auxin efflux transporters PIN2 and PILS6 mediate auxin transport and local accumulation at the root, which in turn triggers developmental response to temperature in the root (Feraru et al., 2019; Hanzawa et al., 2013). Furthermore, the auxin receptors TIR1 and AFB2 are stabilized upon increased ambient temperature by forming a protein complex with HEAT SHOCK PROTEIN 90 (HSP90) and its co-chaperone SUPPRESSOR OF G2 ALLELE SKP1 (SGT1). The accumulation of TIR1 and AFB2 subsequently activates auxin signaling and mediates root thermo-sensory elongation (Wang et al., 2016).

Although root and shoot thermomorphogenesis occur simultaneously during early seedling development (Bellstaedt et al., 2019), it is still unclear whether these responses are coordinated at the whole plant level. In this study, we leveraged a genetic approach combined with comprehensive phenotypic analyses, transcriptional profiling and metabolic measurements to further characterize the molecular circuits mediating root thermomorphogenesis. We found that a shoot regulatory module including HY5, phytochromes and PIF factors can also regulate the root growth response upon perception of higher ambient temperature, demonstrating that shoot and root growth responses are coupled during early seedling development. Furthermore, we show that an additional regulatory axis composed of auxin biosynthesis and perception genes is required during root thermomorphogenesis and propose that the relative abundance of auxin and its downstream signaling activity in the shoot and in the root are critical to coordinately control growth response to temperature in these organs.

## Results

### *HY5* controls the root thermo-response

The impact of increased temperature on plant development has been extensively investigated (reviewed in (Quint et al., 2016)), however it is still unclear whether a core regulatory network governs temperature sensing and signalling in multiple developmental contexts and whether these responses are coordinated across multiple organs. To assess how ambient temperature modulates root development, we grew plants at 21 degree Celsius (°C) and analyzed their growth until three days after transfer at either 21°C or 27°C. In line with previous reports (Feraru et al., 2019; Martins et al., 2017; Wang et al., 2016), wild type plants grown at 27°C displayed an increased primary root growth rate compared to plants kept at 21°C (Figure 1A,B). Having established this experimental set up to analyze root response to temperature shifts, we went on to further characterize the genetic mechanisms underlying this process.

**Figure 1:**
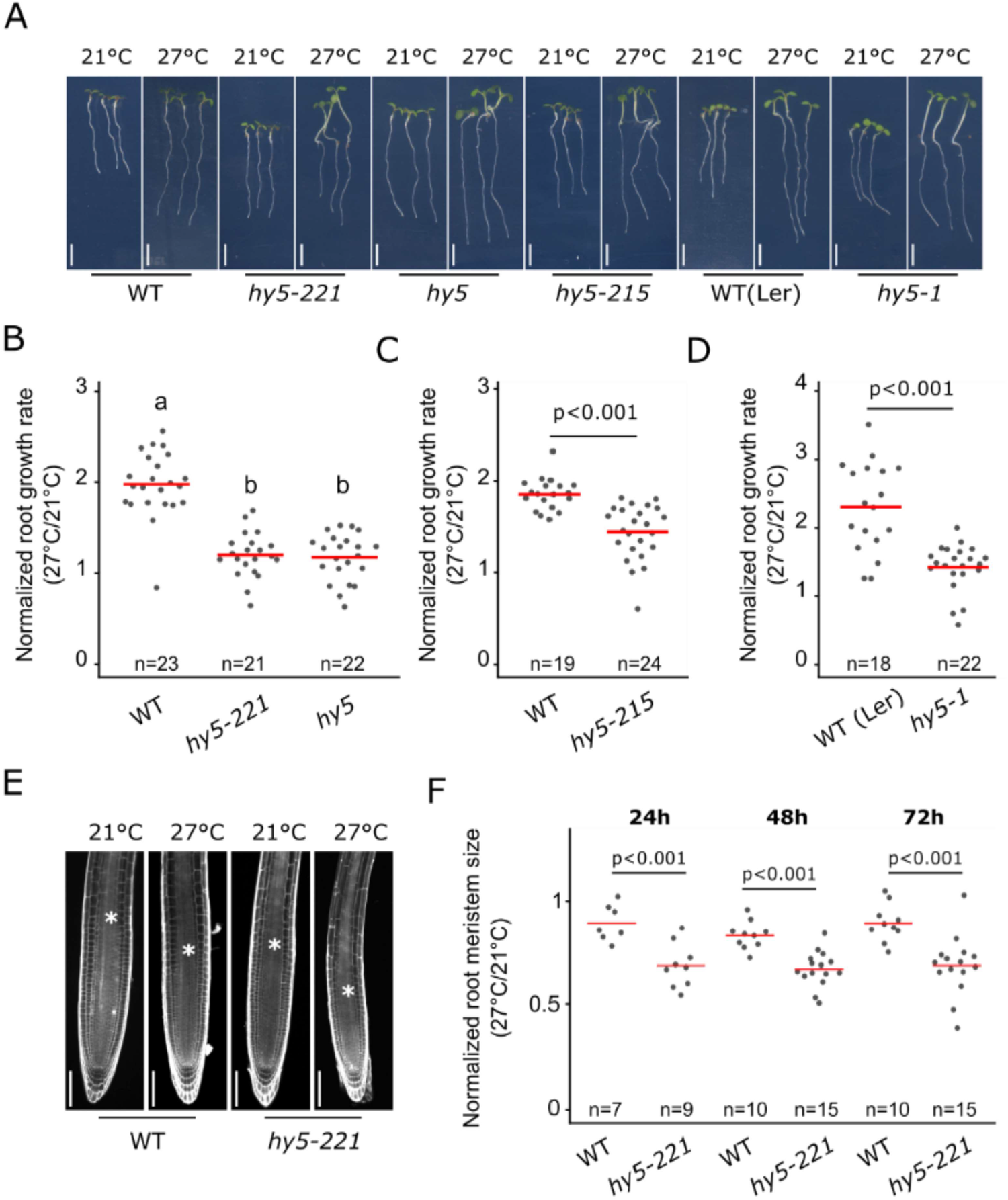
HY5 mediates the root response to higher ambient temperature. (A) Wild type and *hy5* allelic mutant seedling plants 6DAG and 3 days after transfer at 21°C or 27°C. (B-D), Normalized root growth rate (27°C/21°C) in wild type, *hy5, hy5-221, hy5-1* and *hy5-215*. (E) Root meristem in wild type and *hy5-221* 5DAG and 2 days after transfer at 21°C or 27°C. Asterisks mark the root transition zone. (F) Normalized root meristem size (27°C/21°C) in wild type and *hy5-22* at 24, 48 and 72 hours after temperature shift. Statistics: One-way ANOVA, Tukey HSD post-hoc test P<0.05 (A). Student’s t-test (C,D,F). Red bar represents the mean (B,C,D,F). Scale bar: 5mm (A), 100µm (E).

The transcription factor HY5 is a key regulator of shoot thermomorphogenesis, while at the same time regulates root development and hormonal signaling pathways (reviewed in (Gangappa and Botto, 2016)). Thus, we hypothesized that HY5 could regulate the root response to increased ambient temperature. We analyzed the relative root growth rate of *hy5* mutant and wild type plants grown at 21°C and 27°C (Figure 1A-D) and in line with our hypothesis, four different allelic versions of *hy5* mutants displayed reduced root growth response to temperature compared to wild type. While wild type plants increased root growth by 80 to 120%, *hy5* mutants displayed an increase of only 20 to 40% (Figure 1A-D). This reduced response was also observed under a different growth condition with reduced light intensity (see material and methods; source data file) as well as when roots were grown in the dark or on medium not supplemented with sucrose (Supplementary Figure 1A-C), indicating that the reduced response observed in *hy5* was not dependent on light or nutrient conditions. To test whether this reduced response was also associated with changes in root apical meristem (RAM) activity, we measured the dynamics of the root meristem size after temperature shift (Figure 1E,F). Interestingly, *hy5* mutants displayed a lower relative RAM size at all time points analyzed –from 24 hours to 72 hours after temperature shift– indicating that their RAM was hypersensitive to increased ambient temperature compared to wild type plants. Together, these data demonstrate that *HY5* is required to mediate root responses to temperature.

While analyzing the root phenotypes of *hy5* mutants, we observed that plants with a lower root growth frequently displayed longer hypocotyls than plants with a higher root growth, suggesting that shoot and root responses to temperature could be functionally connected. To test this observation, we simultaneously measured hypocotyl and root growth on individual plants and calculated the relative hypocotyl or root growth rate. Raising ambient temperature strongly promoted hypocotyl growth while decreasing root growth response in the *hy5* mutant (Supplementary Figure 1D), supporting the idea that these two processes could be coordinated during early seedling development.

### Phytochromes and PIF activity regulate the root response to higher ambient temperature

The phenotypic relation between hypocotyl and root growth response in the *hy5* mutant suggested that additional regulators of shoot thermomorphogenesis might also modulate the root growth response. Previous studies had demonstrated a critical role of PHYB to sense temperature in the shoot and to mediate hypocotyl growth (Jung et al., 2016; Legris et al., 2016), leading us to hypothesize that the phytochromes might also regulate root thermo-responses. Accordingly, both *phyA* and *phyB* single mutant plants displayed a reduction in the root response to temperature compared to wild type. This difference was further enhanced in *phyAB* double mutants, showing that PHYA and PHYB co-regulate this process (Figure 2A,B). The lower root growth rate in *phyAB* was also associated with a decreased relative root meristem size, demonstrating that root meristematic activity was hypersensitive to increased ambient temperature, similarly to what we observed in *hy5* mutant plants (Figure 2C,D). Collectively, these data demonstrate that in addition to their function in the shoot, the phytochromes are also required for root thermomorphogenesis.

**Figure 2:**
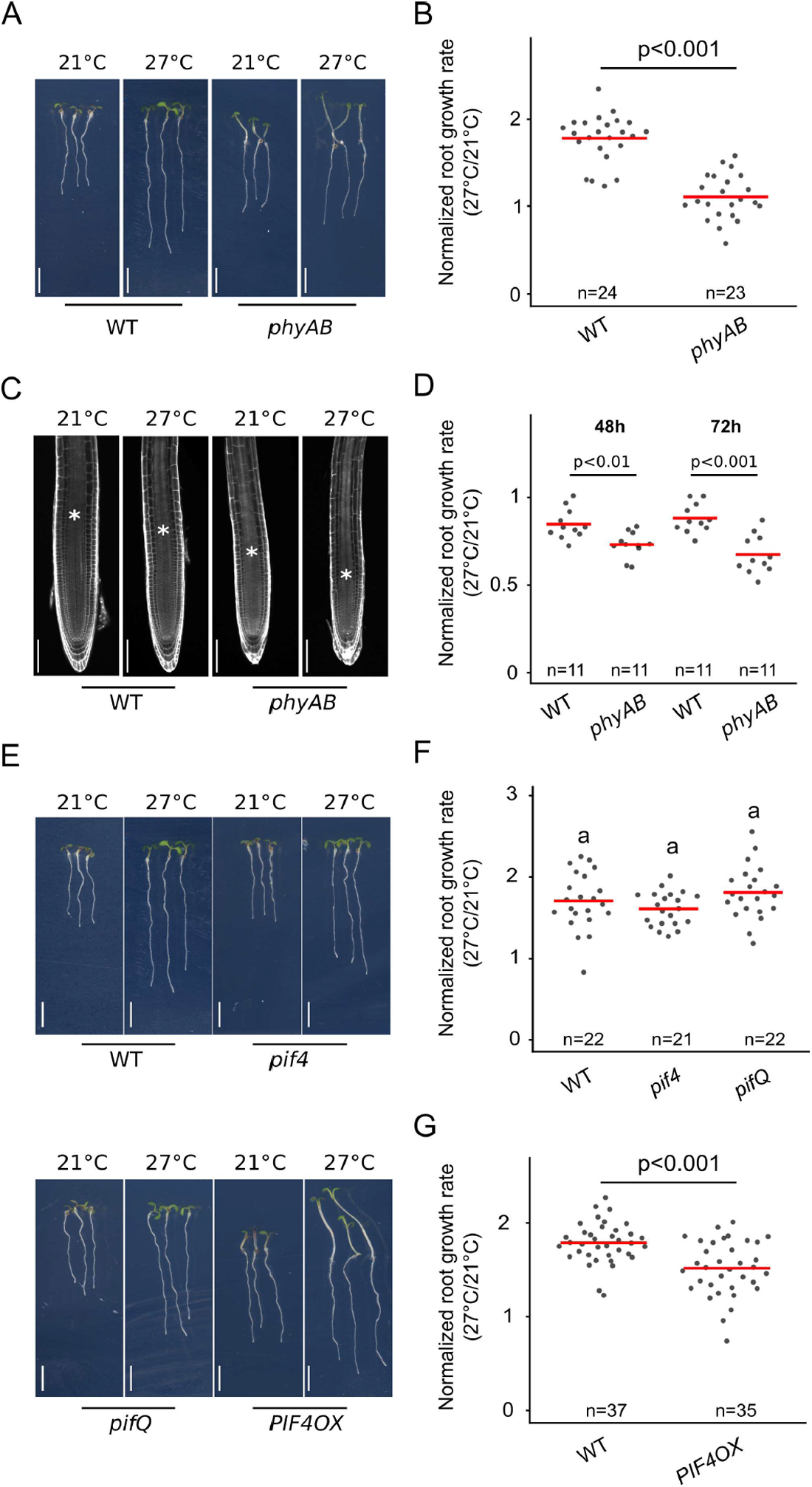
Phytochrome signaling regulates the root response to higher ambient temperature. (A) Wild type (WT) and *phyAB* mutant seedlings 6DAG and 3 days after transfer at 21°C or 27°C. (B) Normalized root growth rate (27°C/21°C) in wild type and *phyAB*. (C) Root meristem in wild type and *phyAB*, 5DAG 2 days after transfer at 21°C or 27°C. Asterisk marks the root transition zone. (D) Normalized root meristem size (27°C/21°C) in wild type and *phyAB*, 48 and 72 hours after temperature shift. (E) Wild type, *pif4, pifQ* and PIF4 OX mutant seedlings 6DAG and 3 days after transfer at 21°C or 27°C. (F-G) Normalized root growth rate (27°C/21°C) in wild type, *pif4, pifQ* (F) and *PIF4 OX* (G).Statistics: One-way ANOVA, Tukey HSD post-hoc test P<0.05 (F). Student t-test (B,D,G). Red bar represents the mean (B,D,F,G). Scale bar: 5mm (A,E), 100µm (C).

Phytochromes mediate the phosphorylation of downstream factors including the PIFs, which are then targeted for degradation (Lorrain et al., 2008). As PIF4 functionally interacts with HY5 during shoot thermomorphogenesis (Delker et al., 2014; Gangappa and Kumar, 2017), we reasoned that PIF4 might also modulate root responses to temperature downstream of the phytochromes. Thus, we tested whether PIF4 and other PIF family members could control root response to temperature. Similarly to previous studies (Martins et al., 2017), *pif4* mutants did not show an impaired root response (Figure 2E,F). Moreover, simultaneously interfering with the function of *PIF1, PIF3, PIF4* and *PIF5* in the *pifQ* mutant had no effect on the root response compared to wild type, indicating that the PIFs were not required to regulate this process (Figure 2E,F). Although the loss-of-function mutants did not display impaired root response to higher temperature, we reasoned that because phytochromes are negative regulators of PIFs, PIF activity might be increased in phytochrome mutants, and that in turn might contribute to the reduction of the root thermo-response in *phyAB* mutants. Thus, we next tested whether promoting PIF function could be sufficient to modulate root growth response. In line with this idea, the gain-of-function *pPIF4:PIF4-FLAG* mutant line (PIF4OX; Gangappa and Kumar, 2017) showed a significant reduction in the root response to higher temperature (Figure 2G), demonstrating that while PIF4 function is not required, it is indeed sufficient to modulate the root response to temperature. As PIF activity is promoted in phytochrome mutants (Park et al., 2018, 2004), our results further suggest that increased PIF4 activity in the *phyAB* could lead to a reduction of the root thermo-response.

### HY5-PIF activity co-regulate root thermomorphogenesis

Having shown that HY5 or phytochromes/PIF activity can modulate shoot and root responses to temperature, we next hypothesized that HY5 and PIFs could co-regulate this process. To test this idea, we first impaired HY5 function together with DET1 and COP1, which are regulators of *PIF4* expression and the hypocotyl response to temperature (Supplementary Figure 2A; Gangappa and Kumar, 2017)). In accordance with a previous report (Gangappa and Kumar, 2017), both *hy5 det1* and *hy5 cop1* double mutants suppressed the enhanced hypocotyl response of *hy5* mutants (Figure 3A,B). Interestingly, these lines also displayed a significant increase in root growth temperature response compared to *hy5* (Figure 3C). These results demonstrated that impairing DET1 and COP1 function can partially rescue root growth rate in response to higher ambient temperature. Importantly, neither *det1* nor *cop1* single mutants displayed an increased root growth response to temperature, suggesting that the genetic interaction between HY5, DET1 or COP1 is critical to modulate root thermomorphogenesis (Supplementary Figure 2B,C). To directly test whether HY5 and PIFs could co-regulate this process, we next simultaneously interfered with HY5 and PIF function using the *hy5 pifQ* quintuple mutant and analyzed growth responses to elevated temperature (Figure 3D-F). Consistent with this idea, both hypocotyl and root growth responses were significantly rescued compared to *hy5* mutants, demonstrating that HY5 and PIF pathways functionally interact to regulate shoot and root responses to temperature (Figure 3D-F). These results demonstrate that the activity of a shoot signaling module including HY5 and PIF genes mediates root response to temperature.

**Figure 3:**
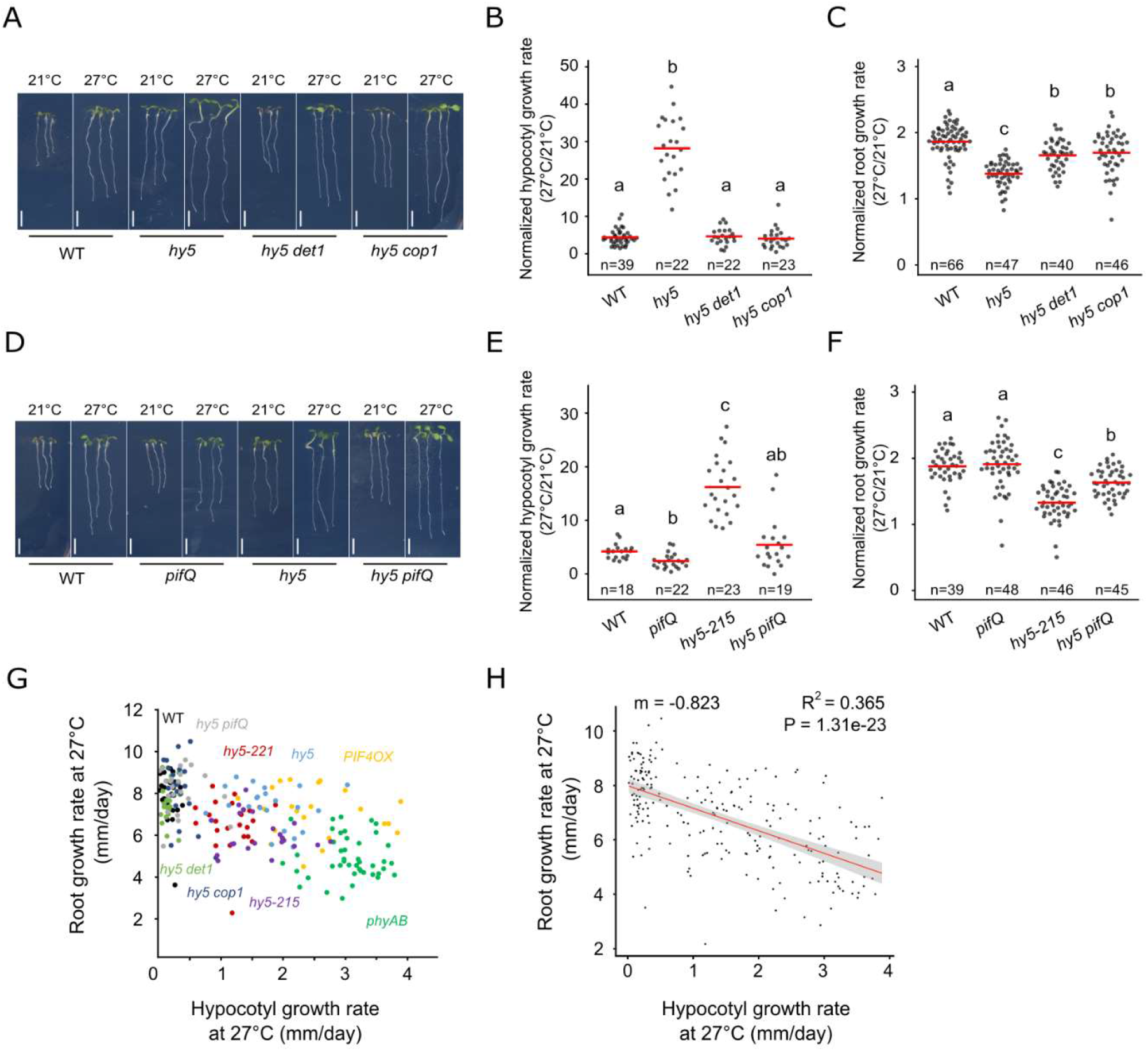
HY5-PIF module regulates the root response to temperature. (A) Wild type, *hy5, hy5 det1* and *hy5 cop1* mutant seedlings 6DAG and 3 days after transfer at 21°C or 27°C. (B-C) Normalized hypocotyl (B) and root growth rate (C) (27°C/21°C) in wild type, *hy5, hy5 det1* and *hy5 cop1*. (D) Wild type, *pifQ, hy5* and *hy5 pifQ* mutant seedlings 6DAG and 3 days after transfer at 21°C or 27°C. (E-F) Normalized hypocotyl (E) and root growth rate (F) (27°C/21°C) in wild type, *pifQ, hy5-215* and *hy5 pifQ*. (G-H) Relation between root and hypocotyl growth rate at 27°C as shown with measurements on individual wild type (n=23), *hy5-221* (n=24), *phyAB* (n=43), PIF4OX (n=22), hy5 (n=22), *hy5 det1* (n=20), *hy5 cop1* (n=22), *hy5-215* (n=23), *hy5 pifQ* (n=22) plants (G) and after non-parametric regression analysis (H). Statistics: One-way ANOVA, Tukey HSD post-hoc test P<0.05 (C,F). One way ANOVA after log10 transformation (B,E), linear regression method, Pearson correlation (H). Red bar represents the mean (B,C,E,F). Scale bar: 5mm (A,D).

Taken together, our phenotypic analyses showed that enhanced shoot growth response was associated with a decreased root response to temperature, further suggesting that shoot and root thermomorphogenesis could be quantitatively negatively correlated. To test this idea, we combined measurements of hypocotyl and root growth of individual plants for nine different genotypes (wild type, *hy5-221, hy5, hy5-215, hy5 pifQ, hy5 cop1, hy5 det1, phyAB* and *PIF4OX*). We then analyzed the relation between hypocotyl and root growth rate at 21°C, 27°C or the relation between their normalized growth rates (Figure 3G; Supplementary Figure 2D-G). Remarkably, we observed that at 27°C, individual genotypes formed distinct groups with root growth rate decreasing as the hypocotyl growth increased, supporting the idea that these traits could be negatively correlated (Figure 3G). We next applied a linear regression model and observed a negative correlation between the root and the hypocotyl growth rate at 27°C (R^2^=0.365), indicating that root growth rate negatively correlates with hypocotyl growth rate at 27°C (Figure 3H). Interestingly, we did not observe this relation at 21°C (R^2^=0.064) or when analyzing temperature responses (R^2^=0.035) (Supplementary Figure 2D-G), indicating that this hypocotyl-root growth correlation is specific to higher ambient temperature conditions. Together, these results show that upon increased ambient temperature, HY5-PIF module is required to balance hypocotyl with root growth responses and further suggest that a developmental trade-off governs hypocotyl and root growth response at higher ambient temperature.

### A shoot to root developmental trade-off in response to higher ambient temperature

The observation that shoot and the root thermomorphogenesis were negatively correlated was intriguing and prompted us to test whether modulating shoot thermo-response was sufficient to impact root growth. To investigate this idea, we used a genetic chimera approach by taking advantage of a HA-YFP-HA-HY5 fusion protein (DOF-HY5) that showed restricted cell-to-cell movement and aimed at driving its expression specifically in the shoot of *hy5* mutants using *CAB3* or *CER6* promoters (Burko et al, 2020b; Procko et al., 2016). In line with previous studies (Chen et al., 2016; Procko et al., 2016), we detected strong accumulation of DOF-HY5 in leaves, petioles and hypocotyls for both constructs, confirming that our *CAB3* and *CER6* promoters were driving expression in the shoot (Supplementary Figure 3A-F; Burko et al, 2020b). While we detected DOF-HY5 accumulation in the root of the *pCER6:DOF-HY5* line, we did not detect fluorescence signal in the root of the *pCAB3:DOF-HY5* lines, indicating that expression driven from the *CAB3* promoter was specific to the shoot and that our tagged version of HY5 was not able to move from the shoot to the root (Figure 4A,C; Supplementary Figure 3A-F). To further confirm these observations, we assessed the accumulation of DOF-HY5 fusion protein either in the root or in the shoot using immunoblotting with a HY5 or a HA antibody (Figure 4D,E). Consistent with our microscopy observations, we observed that DOF-HY5 protein accumulated in the shoot of *pCAB3:DOF-HY5* lines whereas the detected protein levels were similar to *hy5* mutant in the root or were accumulating ubiquitously in the *pCER6:DOF-HY5* line (Figure 4D,E). This provided us with valuable genetic material to further test whether HY5 local activity in the shoot could regulate the root response to temperature.

**Figure 4:**
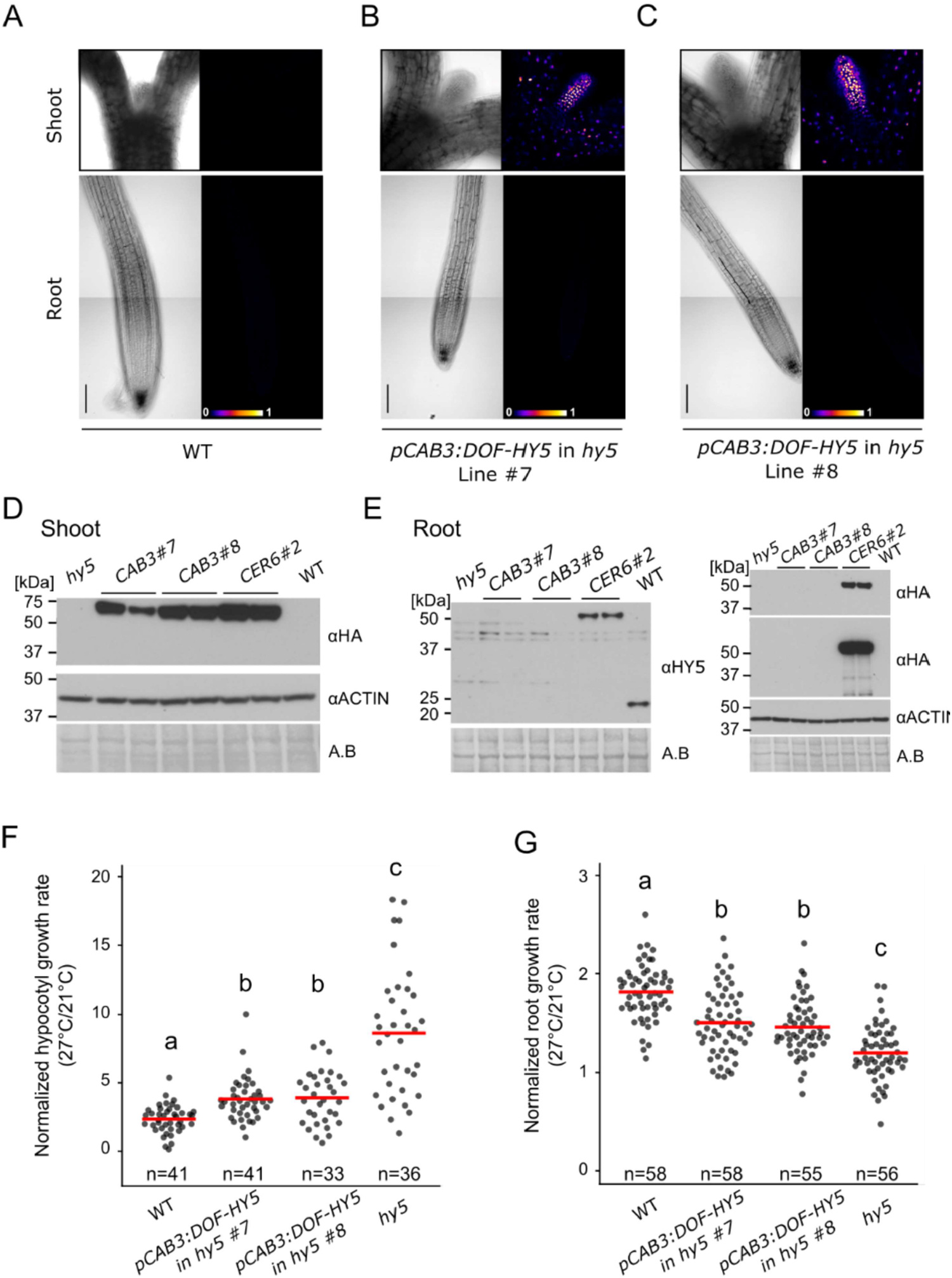
Shoot response to temperature is sufficient to modulate root growth response. (A-C) Brightfield or false color view of wild type seedlings 6DAG (A) and two independent lines of *hy5* carrying *pCAB3:DOF-HY5* (B,C). (D-E) Immunoblotting of shoot (D) or root tissues (E) in wild type (WT), *hy5*, two independent lines of *hy5* carrying *pCAB3:DOF-HY5, hy5* carrying *pCER6:DOF-HY5* and *pCAB3:DOF-HY5* lines at 27°C. DOF-HY5 protein was detected using HA or HY5 antibodies. Amido black staining and actin antibody were used as controls. (F) Normalized hypocotyl growth rate (27°C/21°C) in wild type, *hy5* and *pCAB3:DOF-HY5* rescue lines. (G) Normalized root growth rate (27°C/21°C) in wild type, *hy5* and *pCAB3:DOF-HY5* recue lines. Statistics. One-way ANOVA, Tukey HSD post-hoc test P<0.05 (F,G). Red bar represents the mean (F,G). Scale bar: 100µm (A-C).

We went on to analyze the functionality of the DOF-HY5 fusion protein by measuring hypocotyl and root growth upon response to increased ambient temperature in the *pCER6:DOF-HY5* line. Although *hy5* displayed an increased relative hypocotyl growth rate and a reduced root growth response, these responses were rescued to levels similar to wild type in the *pCER6:DOF-HY5* line, demonstrating that the DOF-HY5 fusion protein was functional (Supplementary Figure 3G-I). These results next prompted us to investigate the local function of HY5 in the shoot during temperature response by analyzing the *pCAB3:DOF-HY5* chimera rescue lines (Figure 4F,G). In line with DOF-HY5 accumulation in the shoot, both *pCAB3:DOF-HY5* lines displayed a partial rescue of the relative hypocotyl growth rate observed in *hy5* (Figure 4F). Strikingly, these two independent lines also showed a significant rescue of the root growth response compared to *hy5*, demonstrating that HY5 function in the shoot was sufficient to modulate root growth response to temperature (Figure 4G). Together, these results reveal that modulating shoot thermomorphogenesis by local HY5 rescue is sufficient to modulate root growth. Together with our previous analyses, these results demonstrate that a developmental trade-off governs hypocotyl and root growth responses to temperature.

### Transcriptional change of metabolic genes in response to temperature

Having shown that a developmental trade-off quantitatively couples shoot and root thermomorphogenesis, we wanted to further delineate the regulatory mechanisms underlying this process. To this end, we used a genome-wide approach and profiled root transcriptomes after a short (4 hours) or a more prolonged (18 hours) temperature treatment using RNAseq.

We first asked whether a core regulatory network could mediate responses to temperature in the root. To strengthen our approach and to alleviate the influence of the genotypes on the response, we compared the transcriptional changes in wild type, *hy5* and *phyAB* plants. Using this method, we identified 327 and 550 genes that were commonly regulated at the early and late time point respectively (Figure 5A,B). Consistent with the temperature treatment imposed onto the plants, the shared regulatory signatures at early time point were associated with heat response (“response to heat”, “response to hydrogen peroxide” and “response to high light intensity”) (Figure 5A). We also observed an enrichment for genes related to metabolism, particularly for members of the glucosinolate biosynthetic pathway and for sucrose transport genes, suggesting that increased ambient temperature might modulate energy metabolism at the root (Figure 5B).

**Figure 5:**
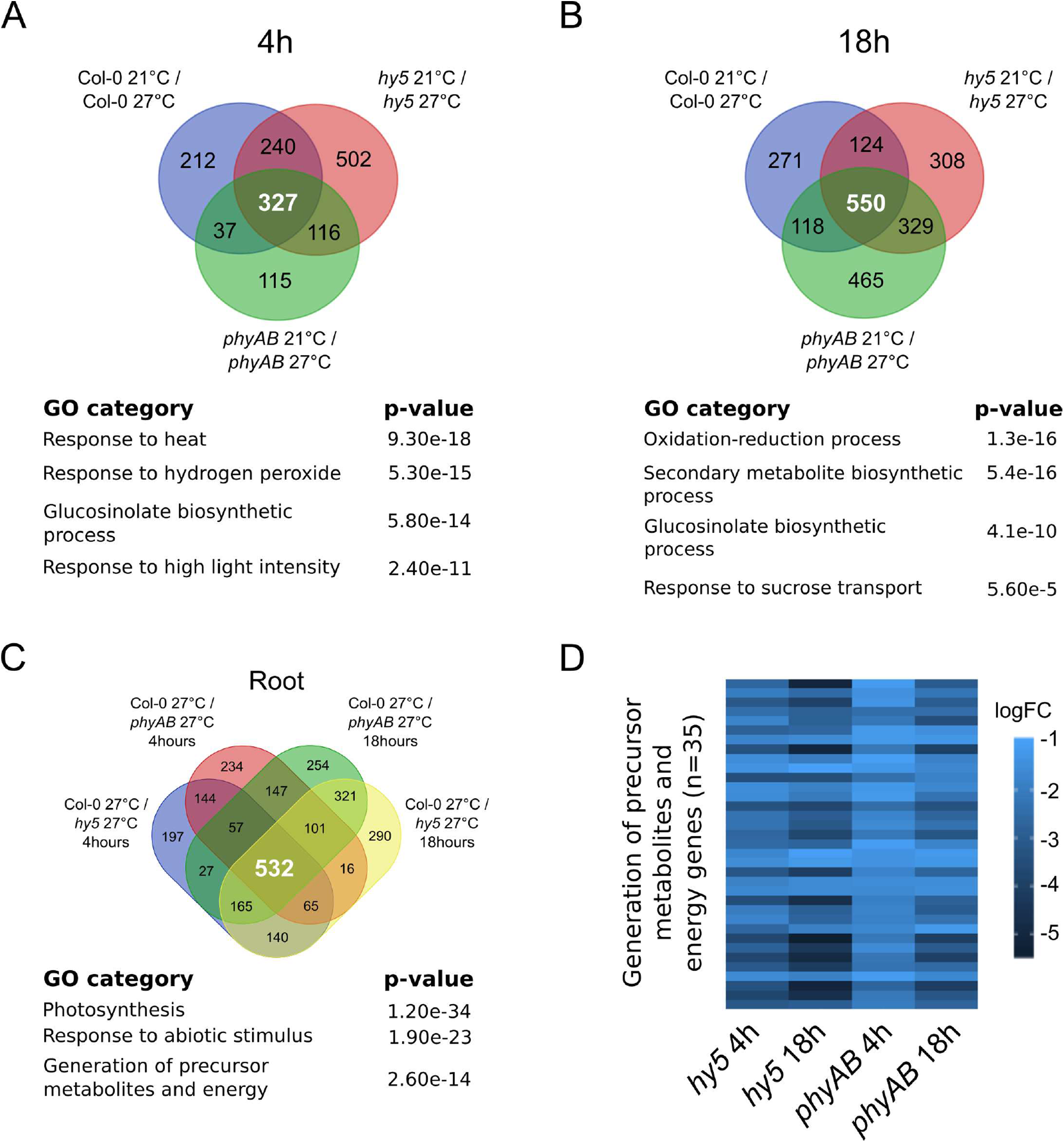
Genome-wide analysis of root response to temperature. (A-B) Genes regulated 4 hours (A) or 18 hours (B) after temperature shift in wild type, *hy5* and *phyAB* roots. Gene ontologies (GO) characterize the biological processes enriched among the temperature-regulated genes that are shared between wild type, *hy5* and *phyAB*. (C) Overlapping misregulated genes in *hy5* and *phyAB* roots at 27°C. (D) Differentially regulated genes belonging to the GO category “Generation of precursor metabolites and energy genes” in *hy5* and *phyAB* roots at 27°C. Statistics: p-value as calculated with AgrigoV2 (A-C).

To further characterize the regulatory function of HY5 and PHYs during root thermomorphogenesis, we next identified genes misregulated in *hy5* and *phyAB* compared to wild type at 27°C (Figure 5C). Strikingly, we observed significant overlap (hypergeometric test; p<0.001) in the sets of genes that were upregulated or downregulated in *hy5* and *phyAB* mutant roots at both time points (Supplementary Figure 4A,B). This overlap supports our previous genetic analyses and demonstrates that HY5 and phytochromes regulate a set of common genes in the root (Figure 5C, Supplementary Figure 4A,B). Among the co-regulated genes, we identified known HY5 target genes – such as *HY5 HOMOLOG* (*HYH*), *SUPPRESSOR OF PHYA* (*SPA*) gene family and *FHY1-LIKE* (*FHL*)– as well as known light signaling genes, which confirmed the quality of our dataset (Supplementary Figure 4C;(Burko et al., 2020; Ciolfi et al., 2013; Lee et al., 2007; Li et al., 2010)). Importantly, we also detected an enrichment for misregulated genes involved in the generation of precursor metabolites, suggesting that the metabolic status was altered in *hy5* and in *phyAB* mutant roots (p=2.6e-14; Figure 5C). Accordingly, all genes belonging to the GO category “generation of precursor metabolites and energy precursor” were significantly downregulated either in *hy5* or *phyAB* at both time points, indicating that HY5 and phytochrome activities are required for the expression of energy metabolism genes in the root (n=35/35, Figure 5D). These results also show that the reduced root growth response observed in *hy5* and *phyAB* mutants correlates with a substantial downregulation of genes involved in the chemical reactions and pathways resulting in the formation of substances from which energy is derived or genes involved in releasing energy from these metabolites. Taken together, the analysis of transcriptional responses suggests that HY5 and phytochrome activity regulate root growth at higher temperature by modulating energy metabolism.

### Auxin perception, signalling and biosynthesis mediate root thermomophogenesis

Having shown that HY5 and phytochromes are required for the expression of energy precursor genes in the root, we next wanted to investigate whether other signals could regulate root thermomorphogenesis downstream of the HY5-PIF module. Some reports have demonstrated that auxin transport and signaling are required for the root response to higher ambient temperature (Feraru et al., 2019; Hanzawa et al., 2013; Wang et al., 2016), however these regulatory interactions have also been challenged and debated (Martins et al., 2017). This prompted us to first confirm the function of auxin homeostasis in our root growth assays.

Shifting plants from 21°C to 27°C led to increased auxin signaling as shown by the increased signal of the *pDR5v2:3xYFP-NLS* transcriptional reporter at the root tip and increased *IAA29* gene expression (Supplementary Figure 5A-C, (Wang et al., 2016)). Consistent with previous reports (Wang et al., 2016), interfering with the auxin receptors TIR1 and AFB2 in *tir1, afb2* and *tir1 afb2* mutant also led to a significant reduction in root growth in response to temperature, demonstrating that auxin perception is required for this response (Figure 6A). To complement these data, we impaired another branch of auxin signaling by interfering with TMKs function, which are membrane localized receptor like kinases involved in the perception of auxin independently of the TIR/AFB system (Cao et al., 2019; Xu et al., 2014). Like observed with the *TIR/AFB* related mutants, we found a reduced root elongation in *tmk1,4* compared to wild type during the root temperature response (Figure 6B). Together, these results confirmed that auxin perception and signaling are required for root thermomorphogenesis.

**Figure 6:**
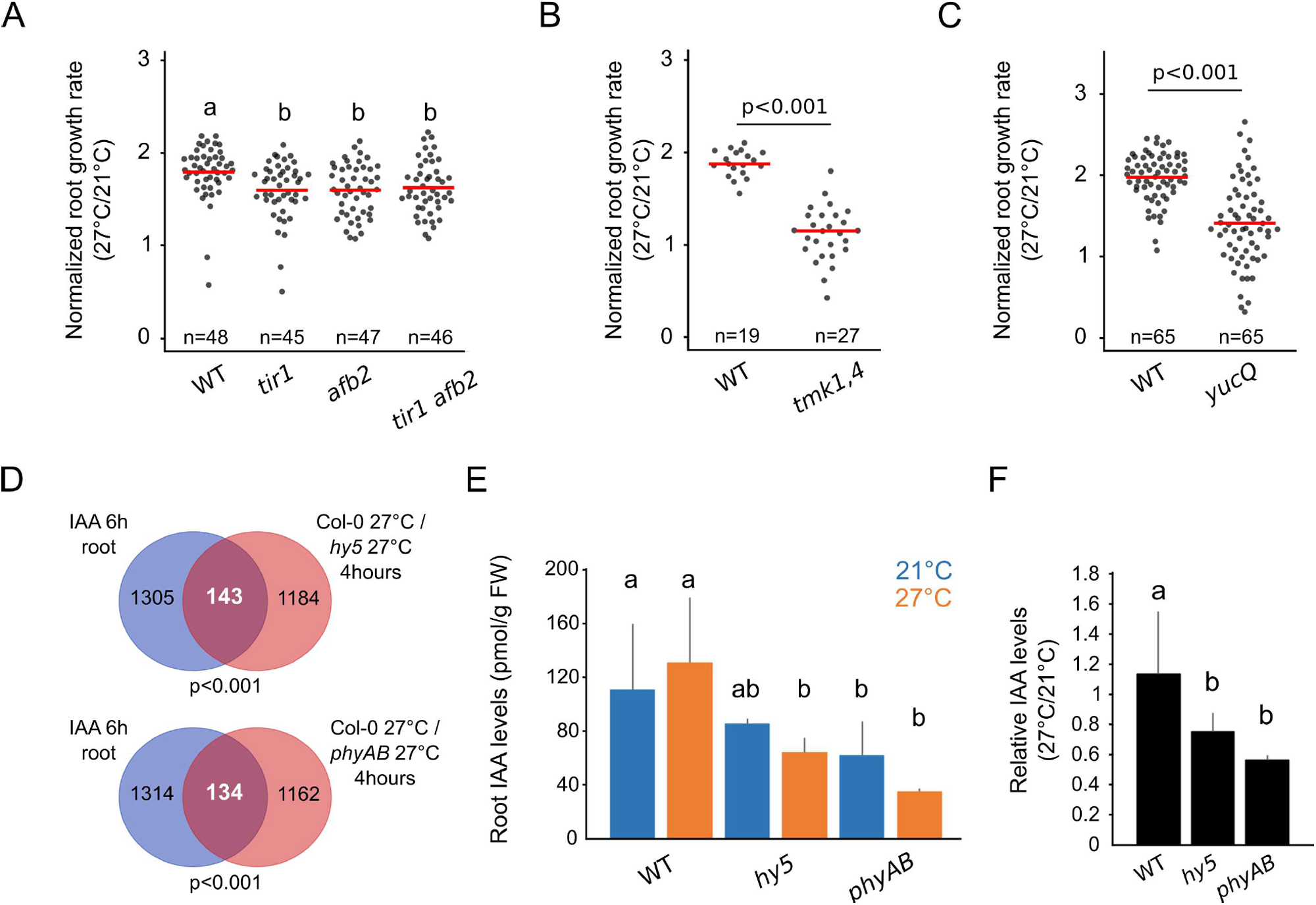
Auxin homeostasis regulates root thermomorphogenesis. (A-C) Normalized root growth rate (27°C/21°C) in wild type, *tir1, afb2, tir1 afb2* (A), *tmk1,4* (B), *yucQ* (C). (D) Differentially regulated genes in *hy5* and *phyAB* roots at 27°C that are auxin responsive according to (Omelyanchuk et al., 2017), 18 hours after temperature shift. (E) IAA concentration (pmol / g of fresh weight (FW)) in roots of seedlings 6DAG, 12 hours after transfer at 21°C or 27°C (n>3). (F) Relative IAA content in root compared to shoot tissues of seedlings 6DAG, 12 hours after transfer at 21°C or 27°C (n>3). Statistics: One-way ANOVA, Tukey HSD post-hoc test p<0.05 (B,D). Student’s t-test (A,C). hypergeometric test (D). One-way ANOVA, Student-Newmann Keuls’s post hoc test p<0.05 (E,F). Red bar represents the mean (A-C).

We next hypothesized that auxin biosynthesis and the control of the hormone level at the root might also modulate root thermomorphogenesis. Thus, we examined the function of auxin biosynthesis by genetically interfering with *YUC* gene activity in the *yuc3,5,7,8,9* (*yucQ*) quintuple mutant. Accordingly, *yucQ* displayed a reduced root growth response compared to wild type, demonstrating that auxin biosynthesis through the activity of the YUCs is also required for root elongation upon higher ambient temperature (Figure 6C). Together, these data unambiguously demonstrate that auxin perception, signaling and biosynthesis are required for root thermomorphogenesis.

### HY5 and phytochromes regulate auxin homeostasis at the root

Having confirmed the function of auxin signaling during root thermomorphogenesis, we next asked whether HY5 and phytochromes could regulate this hormonal pathway. In our root transcriptome, we analyzed the overlap between genes misregulated in *hy5* or *phyAB* at 27°C and auxin responsive genes in the root as obtained from RNA-seq after 6 hours of indole 3-acetic acid (IAA) treatment (Omelyanchuk et al., 2017). Interestingly, we found a significant overlap of genes that were transcriptionally responding to IAA treatment and misregulated in *hy5* and *phyAB* at both early and late time points (Figure 6D, Supplementary Figure 6E). We also found that a significant proportion of genes whose transcriptional response to temperature was differentially regulated in *hy5* or *phyAB* mutants were also responding to auxin in the root (Supplementary Figure 6F). Together these results demonstrated that HY5 and phytochromes activities converged with the auxin regulatory network and further suggested that these factors might control auxin homeostasis during root thermomorphogenesis.

To further examine this idea, we assessed the state of the auxin metabolic pathway by measuring the concentration of IAA and its precursors in roots of wild type, *hy5* and *phyAB* mutants 12 hours after a temperature shift. Surprisingly, we did not observe an increase in total auxin level after temperature shift in wild type root, suggesting that the increase in total auxin level is not required for root thermomorphogenesis, unlike what has been reported for the shoot (Figure 5E;(Gray et al., 1998)). Interestingly, we observed a significant decrease in IAA levels as well as some of the auxin precursors in *hy5* and *phyAB* roots compared to wild type demonstrating that HY5 and phytochromes are required to maintain IAA levels in the root independently of temperature (Figure 6E, Supplementary Figure 5G-H). We also observed a decrease in the relative IAA level in *hy5* and *phyAB* mutants compared to wild type upon increased ambient temperature, indicating that the dynamics of auxin accumulation in the root is impaired upon loss of HY5 and phytochrome activity (Figure 6F). Together these results show that HY5 and phytochrome are required to maintain auxin levels and control the response upon increased temperature in the root. Together, these results also suggest that HY5 and phytochromes control root thermomorphogenesis by regulating auxin levels.

## Discussion

In this study, we investigated the regulatory mechanisms controlling root thermomorphogenesis. Using a genetic approach combined with phenotypic analyses, we find that a regulatory module –including HY5 and phytochromes– concomitantly modulates shoot and root growth responses to higher temperature. In addition, we gain insight on the function of auxin signaling pathway and its connection with HY5 and phytochromes during root thermomorphogenesis (Figure 7). Together, our findings highlight that a developmental trade-off governs shoot and root growth responses and further suggests that roots integrate energy signals with hormonal inputs during thermomorphogenesis.

**Figure 7:**
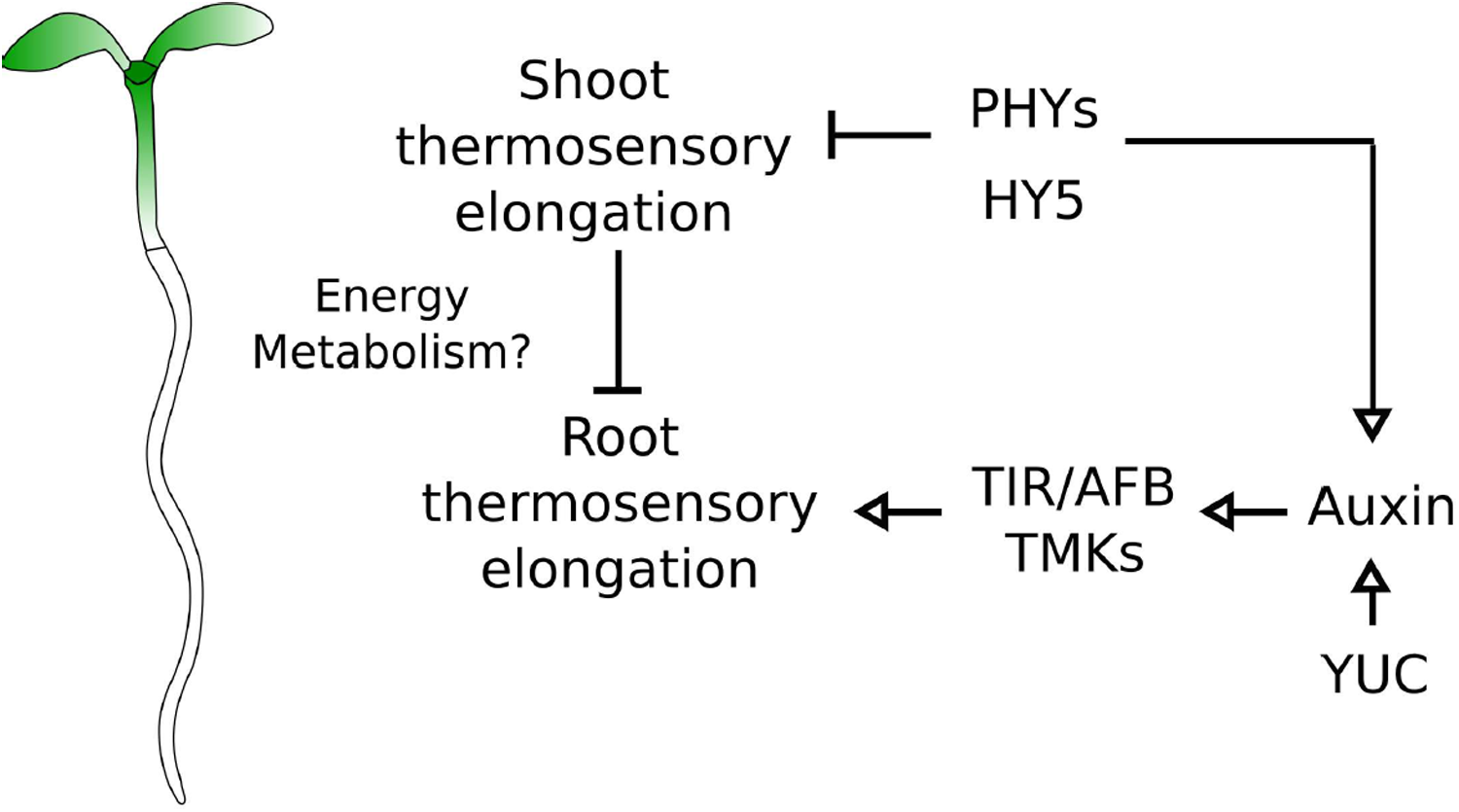
A genetic model for organ growth coordination during plant thermomorphogenesis. Model of root thermosensory response. Roots integrate regulatory signals coming from the shoot through the activity of phytochromes and HY5 with auxin signals mediated by biosynthetic genes (YUC) and signaling (TIR, AFB, TMK).

We showed that HY5 and the phytochromes are required for the root response to temperature. In line with published reports, interfering with PIF activity did not lead to impaired root growth responses to temperature, which previously led to the conclusion that PIFs were not regulating root responses to temperature (Martins et al., 2017). However, we observe that a PIF4 gain-of-function phenocopies the *hy5* and *phyAB* mutant phenotypes, showing that PIF4 is sufficient to regulate root thermomorphogenesis. This result fits well with previous reports showing that PIF activity increases in phytochrome mutants (Park et al., 2018). Furthermore, HY5 acts antagonistically to PIF4 at the promoter of multiple target genes and interfering with HY5 function could enhance PIF4-mediated gene regulation (Gangappa and Kumar, 2017). Accordingly, shoot and root phenotypes in *hy5* mutants are suppressed by dampening PIF expression, demonstrating that HY5 genetically interacts with PIFs during shoot and root thermomorphogenesis. Thus, our results support a model where PIF4 acts downstream of the phytochromes and functionally converges with HY5 to regulate root thermomorphogenesis. Future experiments interfering with phytochromes and PIFs function in higher order mutants will be important to further dissect the function of this regulatory circuit during thermomorphogenesis.

In this context, HY5 also genetically interacts with COP1 and DET1 as shown by the suppression of *hy5* phenotypes in *hy5 det1* and *hy5 cop1*. Interestingly, although the *det1-1* mutant responds similarly to control plants, *cop1-4* shows decreased root growth in response to temperature. This result is intriguing since DET1 and COP1 act together in order to promote HY5 degradation (reviewed in (Lau and Deng, 2012)). Thus, our results also suggest that COP1 can signal independently from HY5 during root thermomorphogenesis. Intriguingly, it was previously shown that COP1 can regulate the polarity of auxin-efflux transporter PIN2 in the root (Sassi et al., 2012). In this context, our findings that *SPA* genes are regulated by HY5 in the root further suggests that the HY5-COP1-SPA module could constitute an intertwined regulatory loop to control root response to temperature. Thus, it will be important to further unravel the function of COP1 or SPAs and to understand whether the interaction with auxin transport is relevant for root thermomorphogenesis.

Our finding that a shoot regulatory module can control hypocotyl growth response and can concomitantly modulate root growth raises interesting questions as to how these two processes are coordinated. Our current data suggest two putative mechanisms that could act in parallel to coordinate shoot with root thermomorphogenesis.

First, our data suggest that temperature responses are tightly connected with the energy metabolism. The observed negative correlation between hypocotyl and root growth responses and the associated downregulation of metabolic precursor genes that play a role in chemical reactions and pathways from which energy is released indicate these two processes could be coordinated by a limitation of metabolic resources that are required during enhanced hypocotyl growth. This hypothesis is consistent with classical studies on biomass allocation between shoots and roots (Shipley and Meziane, 2002; Thornley, 1972). In this context, one possible relevant energy signal could be sucrose, which is produced in the shoot through photosynthesis and has been shown to act as a long-distance signal to promote root growth (Kircher and Schopfer, 2012). Interestingly we found in our genome-wide expression analysis of root responses to temperature that a significant proportion of genes involved in sucrose transport was enriched, suggesting that changes in sugar availability could regulate shoot-to-root growth coordination upon increased ambient temperature. In addition, HY5-PIF4 have been shown to directly regulate the expression of photosynthetic genes and consequently the production of chlorophyll content in young seedlings. Accordingly, *hy5* mutants display lower chlorophyll content than wild type at 27°C (Toledo-Ortiz et al., 2014), which could have a direct impact on the production of photosynthesis-derived sucrose and consequently on root growth. Given that *hy5* mutant still displayed a reduced root response to increased ambient temperature in medium that was not supplemented with sucrose, we believe that external sucrose would have limited impact on this process. Together these data suggest that the availability of energy signals at the shoot could modulate root growth response upon increased ambient temperature.

In parallel to this pathway, another important signal could be the phytohormone auxin. We confirmed that auxin perception and signaling are required for root thermomorphogenesis. Remarkably, we also show that auxin biosynthesis is required, suggesting that the control of auxin levels is critical to regulate root response to temperature. Auxin signaling output is tightly connected to its transport within and across tissues (reviewed in (Benjamins and Scheres, 2008). During shoot responses to temperature, auxin is produced in the cotyledons and transported to the hypocotyl to promote cell elongation (Bellstaedt et al., 2019). Furthermore, the modulation of auxin long-distance transport from the shoot to the root can regulate root developmental responses to environmental light conditions (Sassi et al., 2012). When seedlings are exposed to darkness, auxin levels and signaling decrease in the shoot. This decrease in the shoot concomitantly leads to auxin depletion in the roots as the amount of the hormone transported from the shoot to the root is reduced, thereby inhibiting root growth (Sassi et al., 2012). Our measurements of auxin levels show that *hy5* and phytochrome mutants display lower auxin levels than wild type and that these levels decrease upon increased ambient temperature. These results suggest that the auxin-driven hypocotyl growth in *hy5* and *phyAB* could shift the auxin balance between the shoot and the root, leading to depletion of auxin levels in the root and consequently lead to reduced root growth response to temperature. In contrast to this scenario, other studies suggested that auxin produced in the shoot may not be transported in the root as promoting auxin biosynthesis in the shoot cannot rescue the defects resulting from the loss of function of the *YUC* auxin biosynthesis genes in the root (Chen et al., 2014). Thus, it will be critical to further investigate the dynamics of auxin production, signaling and transport as well as to how it is coordinated between shoot and root during thermomorphogenesis.

Together with our findings, these hypotheses open new avenues to further characterize the communication between shoot and root, which could have important implications for plant growth and biomass allocation upon environmental challenges. Studies have commonly used micro-grafting experiments to investigate long distance signaling between the shoot and the root (Chen et al., 2006, 2016). Given that we analyzed growth response to temperature at early seedling stage, this strategy remains technically challenging as the impact of sectioning on the growth response might override the effect of the genetic backgrounds used as scion. Instead we have used domain-specific rescue approach (Hacham et al., 2011; Kang et al., 2017) by driving a tagged version of HY5 under a shoot-specific promoter. In line with the specificity of the shoot expression, we did not detect fluorescent signal, nor HY5 protein accumulation in the root by immuno-blotting. Although these experimental methods cannot fully exclude that traces of HY5 protein are still present, the levels would be considerably lower than the wild type and unlikely to have strong impact on the observed phenotype. Thus, we propose that using shoot-specific or root-specific genetics with tools such as the CAB3 or the INORGANIC PHOSPHATE TRANSPORTER 1-1 (PHT1:1) promoters (Procko et al., 2016; Vijaybhaskar et al., 2008) could be valuable to further elucidate how the shoot and the root communicate during thermomorphogenesis.

Based on our results, we propose a model where roots integrate systemic signals modulated by a shoot module including HY5, phytochromes with more locally acting auxin signaling during thermomorphogenesis (Figure 7). The integration of signals that are relayed from the shoot as well as more local ones in the root could constitute a flexible system to adapt growth in response to changes in air temperature perceived in the shoot while at the same time tuning growth locally by modulating hormonal homeostasis. Thus, it will be important in the future to further understand to what extent these two signaling pathways interact and how they are coupled at the temporal level.

## Material and methods

### Plant material and growth conditions

In this study we used the following published lines: *phyAB* (Zheng et al., 2013), *hy5* (Jia et al., 2014), *hy5-221, hy5-215, hy5-1* (Oyama et al., 1997), *phyA-211* (Reed et al., 1994), *phyB-9* (Reed et al., 1993), *pif4-101* (Lorrain et al., 2008), *pif1,3,4,5* (*pifQ*)(Leivar et al., 2008), PIF4-OX (pPIF4:PIF4-FLAG)(Gangappa and Kumar, 2017), *hy5 pifQ* (Jia et al., 2014), det1-1 (Pepper et al., 1994), cop1-4 (McNellis et al., 1994), *hy5 det1* (Gangappa and Kumar, 2017), *hy5 cop1* (Rolauffs et al., 2012), *tir1-1, afb2-3, tir afb2* (Parry et al., 2009), *tmk1 tmk4* (Dai et al., 2013), *yucca3,5,7,8,9* (*yucQ*) (Chen et al., 2014), DR5v2 (Liao et al., 2015). CAB3, CER6 promoters were previously described (Procko et al. 2016) and the DOF (HA-YFP-HA) tag was described in (Burger et al. 2017). HY5 rescue lines were generated by inserting *pCAB3:HA-YFP-HA-HY5* and *pCER6:HA-YFP-HA-HY5* in the *hy5* background (Lian et al., 2011) as described in (Burko et al, 2020b).

When not specified, plants were grown in long day conditions (16/8h) in walk-in growth chambers (Conviron, Winnipeg, Manitoba, Canada) at 21°C or 27°C, 60% humidity, at 146 PAR (see source data for light spectra). During nighttime, temperature was decreased to 15°C and 21°C respectively. In our growth condition 2, plants were grown in reach-in growth chambers at 60% humidity, 122 PAR (see source data for light spectra), temperature was kept constant at either 21°C or 27°C. Environmental conditions were established and monitored with commercial software (Valoya, Helsinki, Finland).

Roots grown on plates in the dark were isolated from light using metal combs that contained holes and plates were wrapped with aluminum foil.

Plants were cultivated on plates containing ½ Murashige Skoog (Caisson, Smithfield, UT, USA), 1%MES (Acros Organic, Hampton, NH, USA), 1% sucrose (Fisher Bioreagents, Hampton, NH, USA) and 0.8% Agar powder (Caisson, Smithfield, UT, USA). For temperature shift experiments, plants were germinated and grown until 3 days after germination at 21°C to synchronize their development. On the third day, plants were shifted at ZT1-3 at 27°C and grown for 3 additional days at 21°C or 27°C.

### Root measurements and analysis

Root images were acquired using a multiplex scanning system as described in (Slovak et al., 2014). Images were processed using the Fiji software (https://fiji.sc/). Root and hypocotyl lengths were measured at 3DAG (before temperature shift) and at 6DAG. Growth rate were obtained by subtracting the length at 6DAG and 3DAG. Normalized growth rates were calculated by dividing root growth rate at 27°C by the average growth rate at 21°C. Raw values for individual temperatures can be found in the source data file.

For the time course analysis of normalized root growth rate, plates were scanned at 0, 12, 24, 48, 72 hours after temperature shift. Images were stacked with Image J and root length was measured at individual time points.

Statistical analysis was performed using Excel (Microsoft, Redmond, WA, USA) or R software (https://www.r-project.org/). Linear regression was performed using the lm function in R and graph displayed with ggplot2 (https://www.r-project.org/) (codes are available upon request).

Confocal pictures were acquired on a Zeiss 710 inverted microscope (Zeiss, Oberkochen, Germany) or on Zeiss CSU Spinning Disk Confocal Microscope (Salk Biophotonics Core). Pictures were processed using Fiji software (https://fiji.sc/). Root meristem size was measured from the quiescent center to the first cortical cell that is twice as long as wide as was previously described (Feraru et al., 2019).

Dot plots were generated using the plots of data online tool (Postma and Goedhart, 2019).

### Immunoblotting

Western blots were performed as described in (Li et al., 2012) with minor modifications. 25 roots and 20 shoots were harvested at 6DAG and extracted in 2X loading buffer (36µl bME+1ml 4x loading buffer). Loading buffer was added to roots (70µl) and shoots (140uµl) and then boiled for 5 min. Bis-tris gel 4-12% (Invitrogen, Carlsbad, CA, USA) and semi-dry transfer (Pierce G2 Fast Blotter, Thermo Scientific, Waltham, MA, USA) were used. Primary antibodies used were αHA-HRP 1:2000 (12013819001 Roche), αHY5(N) 1:5000 (R1245-1b ABicode), αActin 1:30,000 (A0408 Sigma).

### Gene expression analysis

Biological triplicates were analysed. Total RNA was extracted from roots or shoot of plants 6 DAG using RNA easy kit (Qiagen, Hilden, Germany). RNA was treated with DNAse using the Turbo DNA-free kit (Invitrogen, Carlsbad, CA, USA) and further purified on columns from the RNA easy kit.

Next generation sequencing (NGS) library was generated using the TruSeq Stranded mRNA library prep kits (Illumina, San Diego, CA, USA). Libraries were sequenced on HiSeq2500 (Illumina, San Diego, CA, USA) as single read 50bases. Raw reads can be found at GEO under the number: GSE138133.

NGS analysis was performed using Tophat2 for mapping reads on the Arabidopsis genome (TAIR10) (Kim et al., 2013, p. 2), HT-seq for counting reads (Anders et al., 2014) and EdgeR for quantifying differential expression (Robinson et al., 2009). We set a threshold for differentially expressed genes (Fold change (FC) >2 or FC<-2, FDR<0.01). Genotype x Environment interaction analysis was performed using linear model and type II ANOVA in R (codes are available upon request).

Gene ontology analysis was performed using AgriGOv2 online tool (Tian et al., 2017). Venn diagrams were generated with the VIB online tool (http://bioinformatics.psb.ugent.be/webtools/Venn/).

### Auxin measurements

For auxin measurement, plants were shifted at ZT1-3 at 27°C, grown at 21°C or 27°C and harvested at ZT 13-15.

The extraction, purification and the LC-MS analysis of endogenous IAA, its precursors and metabolites were carried out according to (Novák et al., 2012). Briefly, approx. 10 mg of frozen material per sample was homogenized using a bead mill (27 hz, 10 min, 4°C; MixerMill, Retsch GmbH, Haan, Germany) and extracted in 1 ml of 50 mM sodium phosphate buffer containing 1% sodium diethyldithiocarbamate and the mixture of ^13^C_6_- or deuterium-labeled internal standards. After centrifugation (14000 RPM, 15 min, 4°C), the supernatant was divided in two aliquots, the first aliquot was derivatized using cysteamine (0.25 M; pH 8; 1h; room temperature; Sigma-Aldrich), the second aliquot was immediately further processed as following. The pH of sample was adjusted to 2.5 by 1 M HCl and applied on preconditioned solid-phase extraction column Oasis HLB (30 mg 1 cc, Waters Inc., Milford, MA, USA). After sample application, the column was rinsed with 2 ml 5% methanol. Compounds of interest were then eluted with 2 ml 80% methanol. Derivatized fraction was purified alike. Mass spectrometry analysis and quantification were performed by an LC-MS/MS system comprising of a 1290 Infinity Binary LC System coupled to a 6490 Triple Quad LC/MS System with Jet Stream and Dual Ion Funnel technologies (Agilent Technologies, Santa Clara, CA, USA).

Raw measurements for individual temperatures can be found in the source data file.

## Competing interests

The authors declare no competing interests

## Acknowledgements

We would like to thank Yvon Jaillais (ENS, Lyon, France), Mark Estelle (UCSD, La Jolla, USA), Yunde Zhao (UCSD, La Jolla, USA), and members of the Chory lab (Bjorn Willige, Adam Seluzicki; Salk Institute, La Jolla, USA) for kindly sharing published mutant plant lines with us. We would also like to thank members of the Busch laboratory for critically reading the manuscript. This study was funded by the National Institute of General Medical Sciences of the National Institutes of Health (grant number R01GM127759 to W.B.) and start-up funds from the Salk Institute for Biological Studies. J.C. is investigator of the Howard Hughes Medical Institute. This study was supported by the HHS NIH National Institute of General Medical Sciences (grant 5R35GM122604-02_05 to J.C.), the Howard Hughes Medical Institute (to J. C.), the European Molecular Biology Organization (grant ALTF 785-2013 to Y.B.), the United States-Israel Binational Agricultural Research and Development Fund (grant FI-488-13 to Y.B). K.L. and J.S. acknowledge the Swedish research councils VINNOVA, VR and the Knut and Alice Wallenberg Foundation (KAW). They also thank the Swedish Metabolomics Centre (http://www.swedishmetabolomicscentre.se/) for access to instrumentation.

---

**Supplementary Figure 1:**
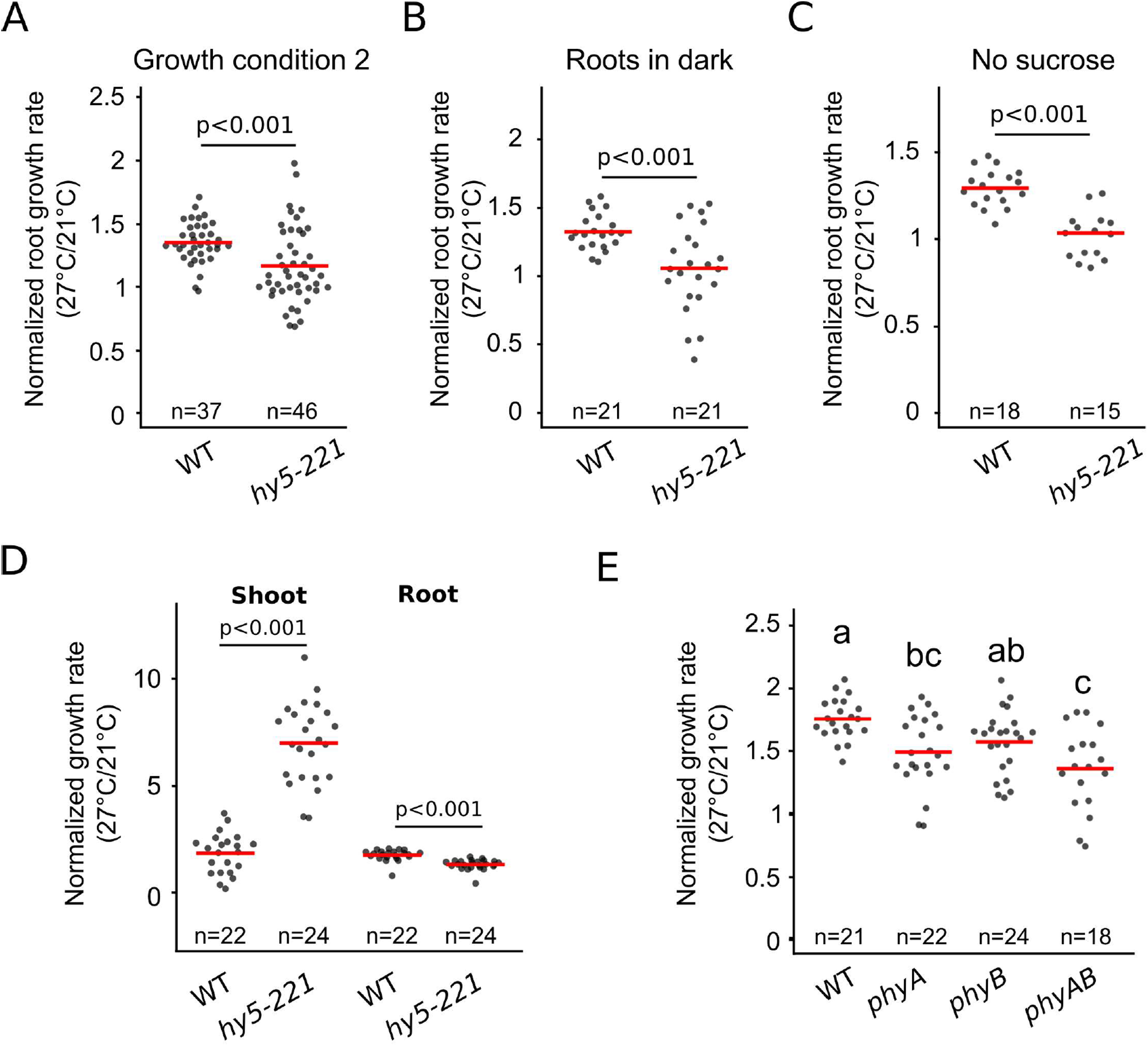
Characterization of *hy5* in response to higher ambient temperature. (A-C) Normalized root growth rate (27°C/21°C) in wild type and *hy5-221* under growth condition 2 (A) (see method section) or with roots grown in the dark (B) or with roots grown on medium not supplemented with sucrose (C). (D) Normalized hypocotyl and root growth rate (27°/21°C) simultaneously measured on individual plants. (E) Normalized root growth rate (27°C/21°C) in wild type, *phyA, phyB* and *phyAB*. Statistics: Student’s t-test (A-D), one-way ANOVA, Tukey HSD post-hoc test P<0.05 (E). Red bar represents the mean (A-E)

**Supplementary Figure 2:**
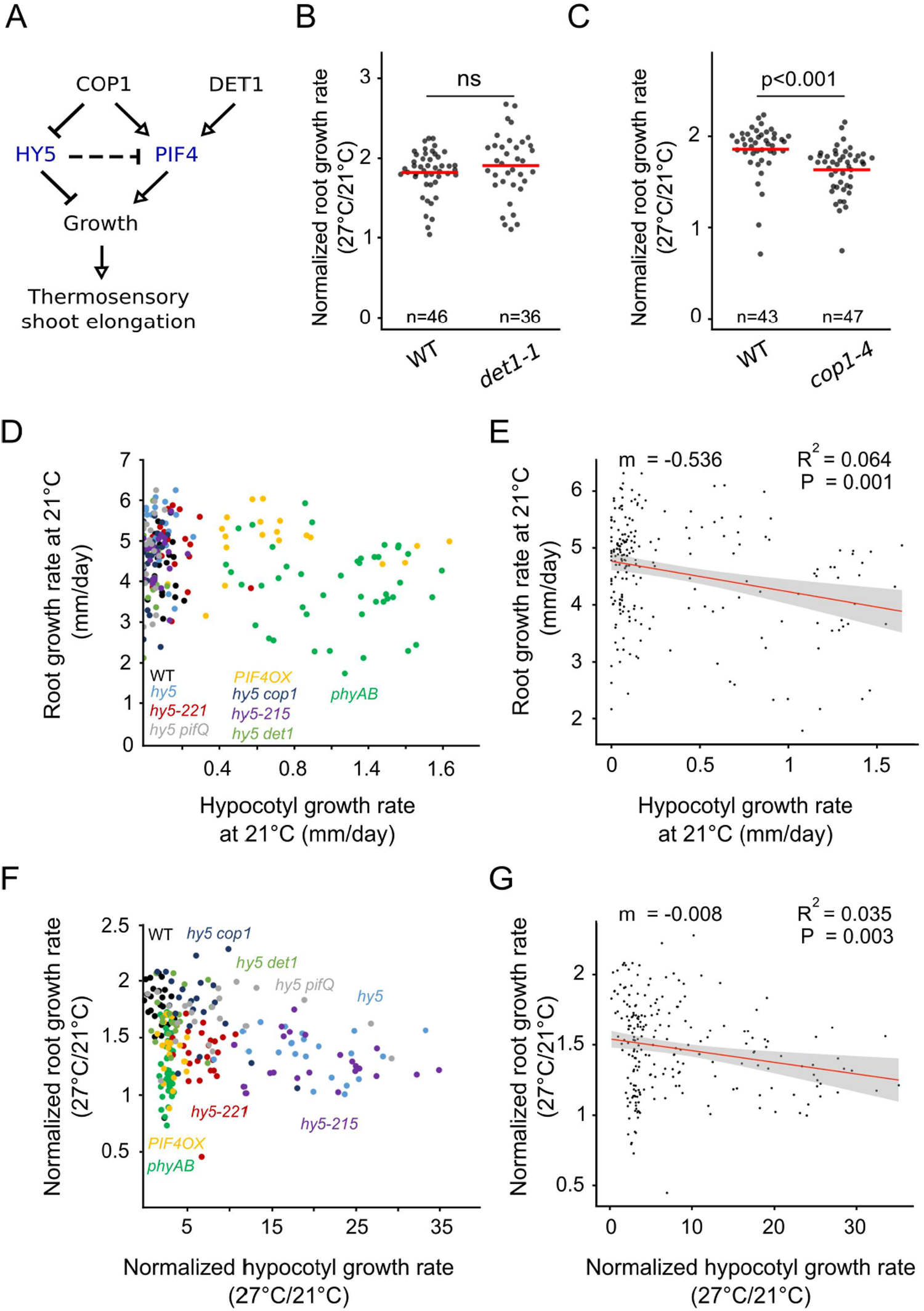
Characterization of the HY5-PIF module in response to higher ambient temperature. (A) Regulatory model of thermosensory shoot elongation as proposed by (Delker et al., 2014). (B-C) Normalized root growth rate (27°C/21°C) in wild type, *det1-1* (B) and *cop1-4* (C). (D-E) Relation between root and hypocotyl growth rate at 21°C as shown with measurements on individual wild type (n=20), *hy5-221* (n=19), *phyAB* (n=44), PIF4OX (n=20), hy5 (n=22), *hy5 det1* (n=19), *hy5 cop1* (n=22), *hy5-215* (n=23), *hy5 pifQ* (n=20) plants (D) and after non-parametric regression analysis (E). (F-G) Relation between normalized root and hypocotyl growth rate (27°C/21°C) as shown with measurements on individual wild type (n=23), *hy5-221* (n=24), *phyAB* (n=43), PIF4OX (n=22), hy5 (n=22), *hy5 det1* (n=20), *hy5 cop1* (n=22), *hy5-215* (n=23), *hy5 pifQ* (n=22) plants (F) and after non-parametric regression analysis (G) Statistics: Student’s t-test (B,C), linear regression method, Pearson correlation (E,G). Red bar represents the mean (B,C).

**Supplementary Figure 3:**
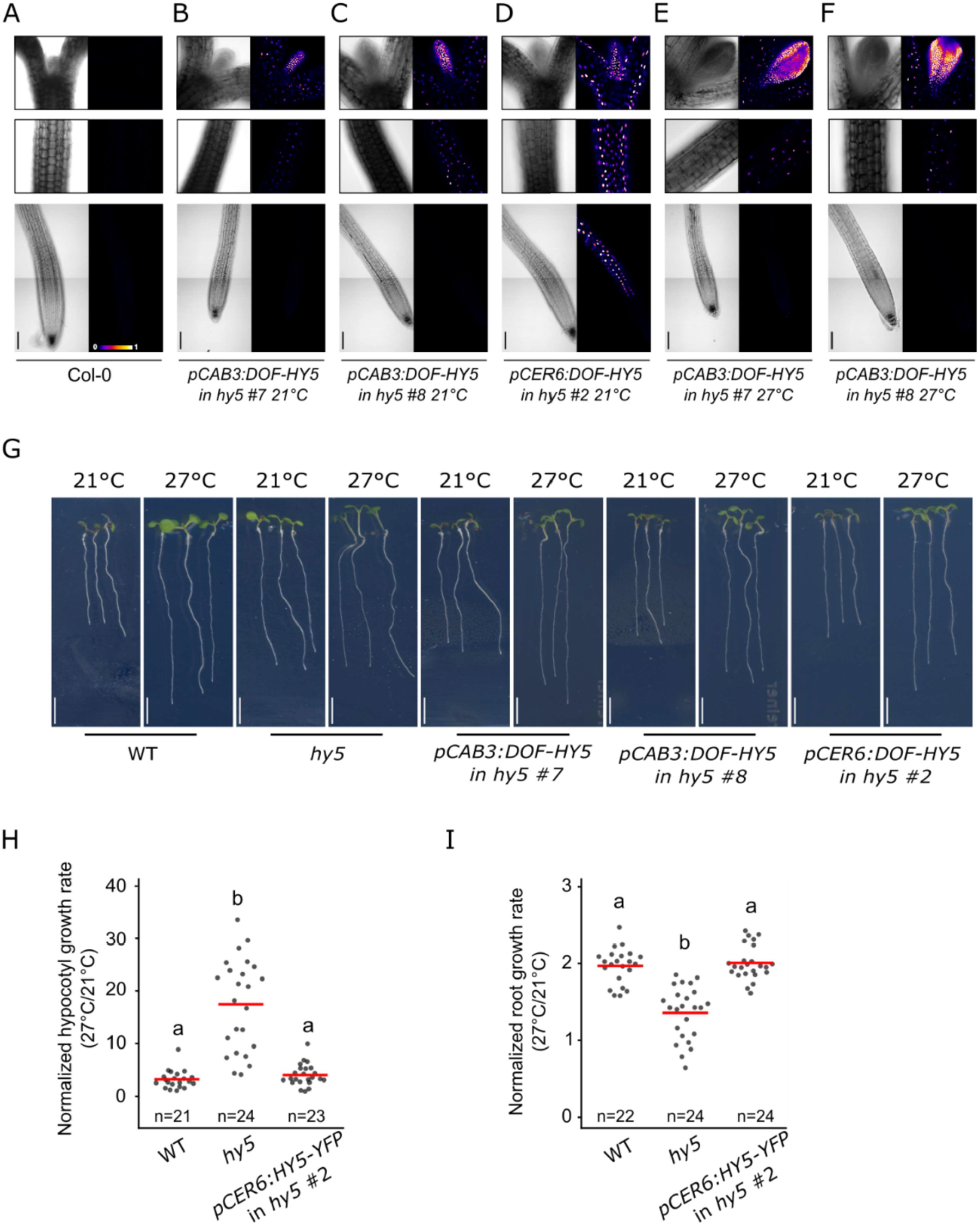
Characterization of HY5 chimera lines. (A-F) Brightfield or false color view of seedlings roots, hypocotyls and apical ends 6DAG in Col-0 (A), two independent lines of *hy5* carrying *pCAB3:DOF-HY5* (B,C), *hy5* carrying *pCER6:DOF-HY5* (D) and *pCAB3:DOF-HY5* lines at 27°C (E,F). (G) Wild type, *hy5*, two independent lines of *hy5* carrying *pCAB3:DOF-HY5, hy5* carrying *pCER6:DOF-HY5* seedlings 6DAG and 3 days after transfer at 21°C or 27°C. (H-I) Normalized hypocotyl (H) and root growth rate (I) (27°C/21°C) in wild type, *hy5* and *hy5* carrying *pCER6:DOF-HY5*. Statistics: One-way ANOVA, Tukey HSD post-hoc test p<0.05 (H,I). Red bar represents the mean (H,I). Scale bar: 5mm (G), 100µm (A-F).

**Supplementary Figure 4:**
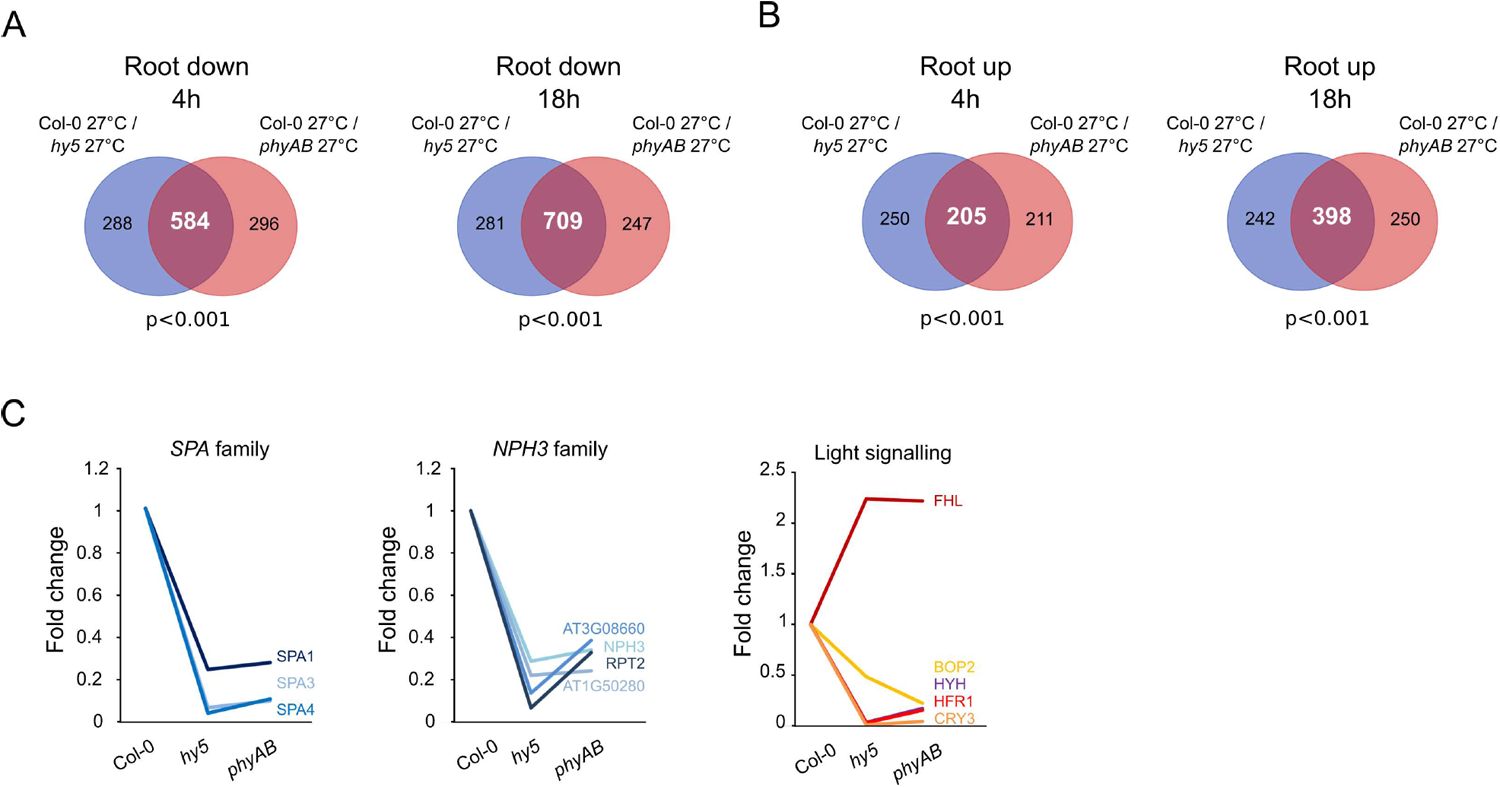
Transcriptional profiling of *hy5* and *phyAB* mutants. (A-B) Commonly downregulated (A) or upregulated (B) genes in *hy5* and *phyAB* roots at 27°C, 4 or 18 hours after temperature shift. (C) Root expression change of genes co-regulated by HY5 and phytochromes.. Statistics: hypergeometric test (A,B)

**Supplementary Figure 5:**
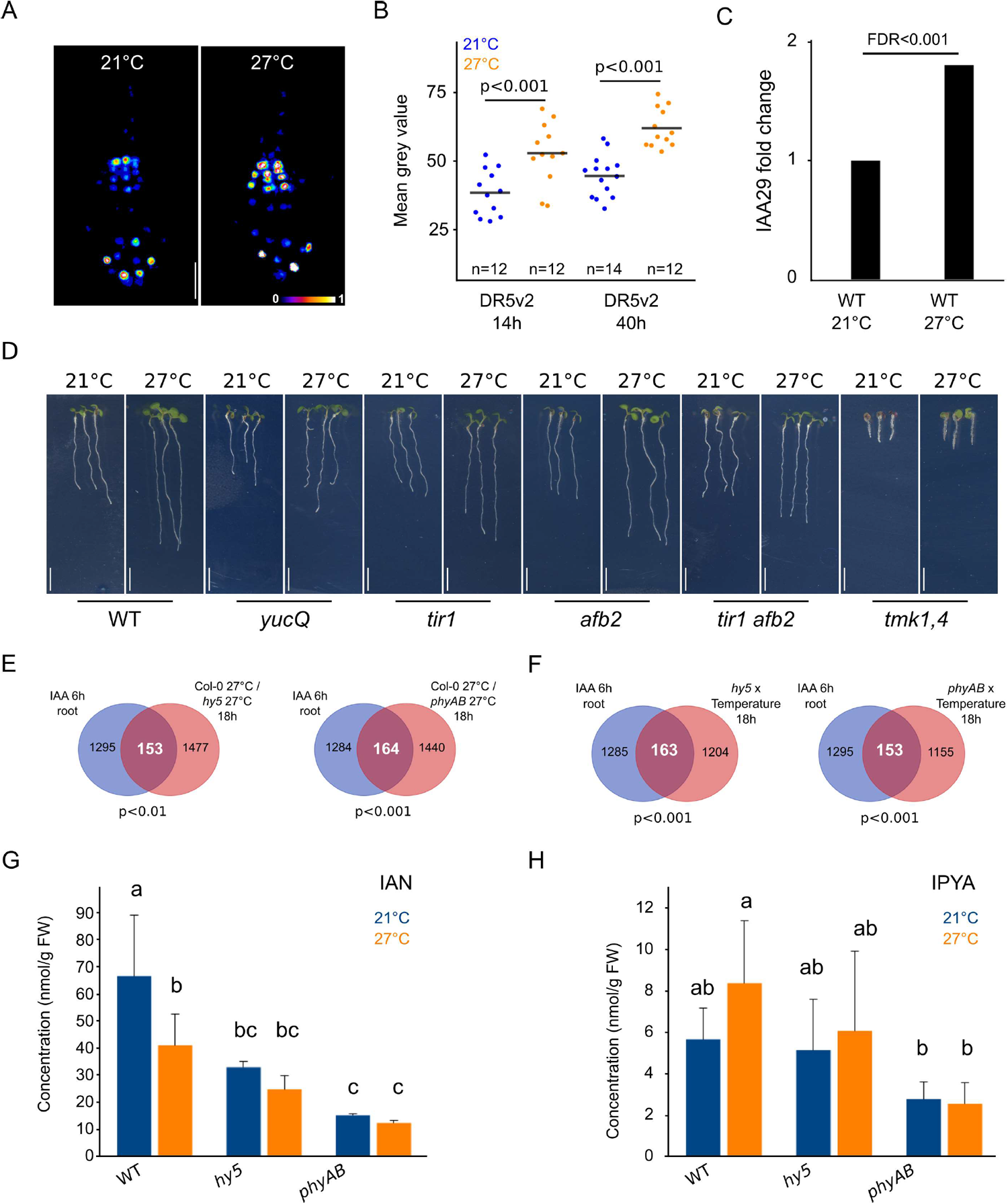
Auxin homeostasis during root thermomorphogenesis. (A) False color view of root tips expressing *pDR5v2:3xYFP-NLS* grown at 21°C or 27°C. (B) Quantification of DR5v2 signal in seedlings 14 hours or 40 hours after transfer at 21°C or 27°C. (C) Relative IAA29 expression detected in wild type roots 4 hours after transfer at 21°C or 27°C. (D) Wild type, *yucQ, tir1, afb2, tir1 afb2* and *tmk1,4* mutant seedlings 6DAG and 3 days after transfer at 21°C or 27°C. (E) Differentially regulated genes in *hy5* and *phyAB* roots at 27°C, 18 hours after temperature shift that are auxin responsive (Omelyanchuk et al., 2017). (F) Overlap between genes whose transcriptional response changes in *hy5* or *phyAB* and that are auxin responsive. (G-H) Indole-3-acetonitrile (G) and indole-3-pyruvic acid (H) concentration in 6DAG wild type, *hy5-221* and *phyAB* roots 12 hours after transfer at 27°C. Statistics: Student t-test (B), false discovery rate (FDR) as calculated by EdgeR (C), hypergeometric test (E-F), one-way ANOVA, Student-Newmann Keuls’s post hoc test p<0.05 (G-H). Scale bar: 5mm (D). Black bar represents the mean (B).

## Supplementary data root measurements

### Statistics

One-way ANOVA, Tukey HSD post-hoc test p<0.05

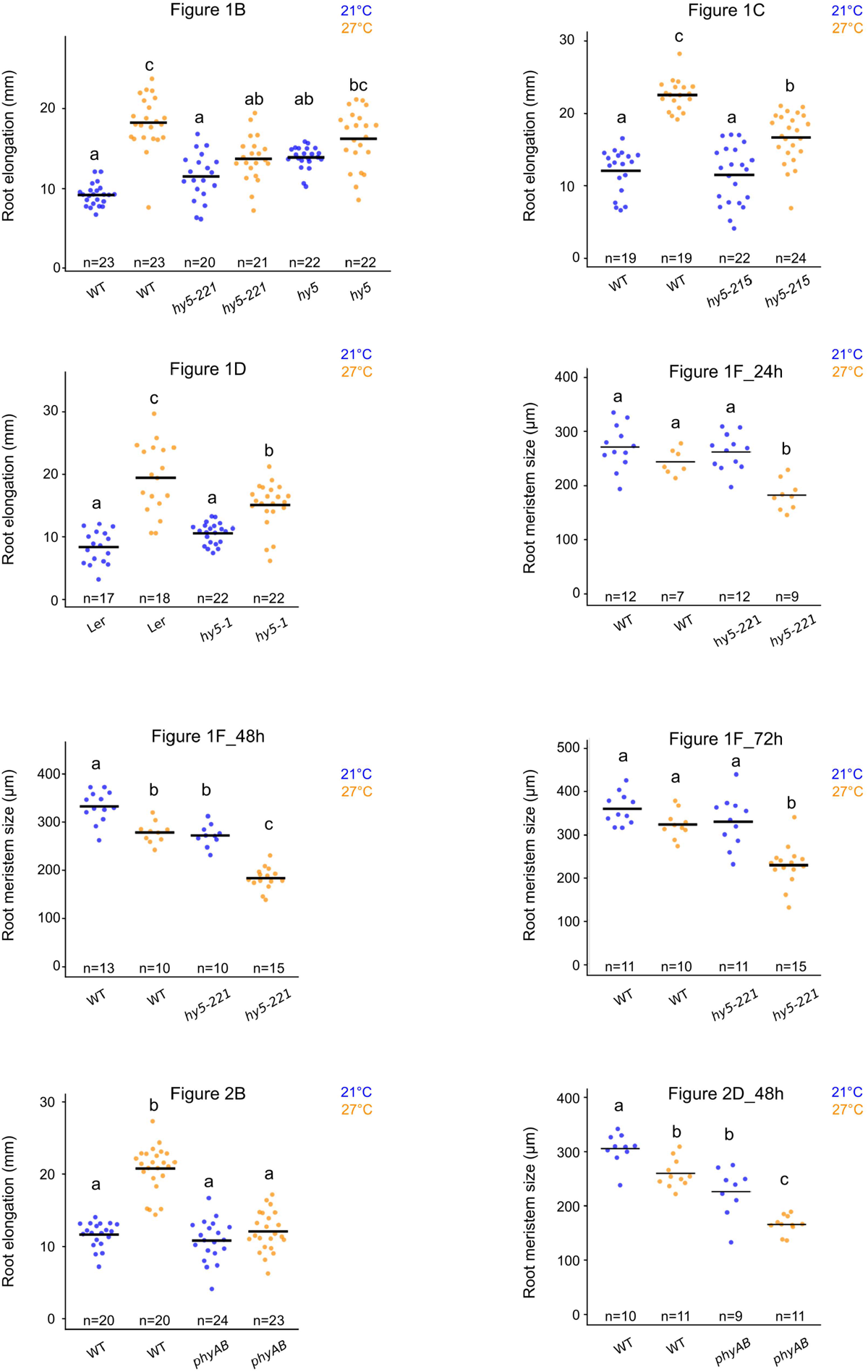

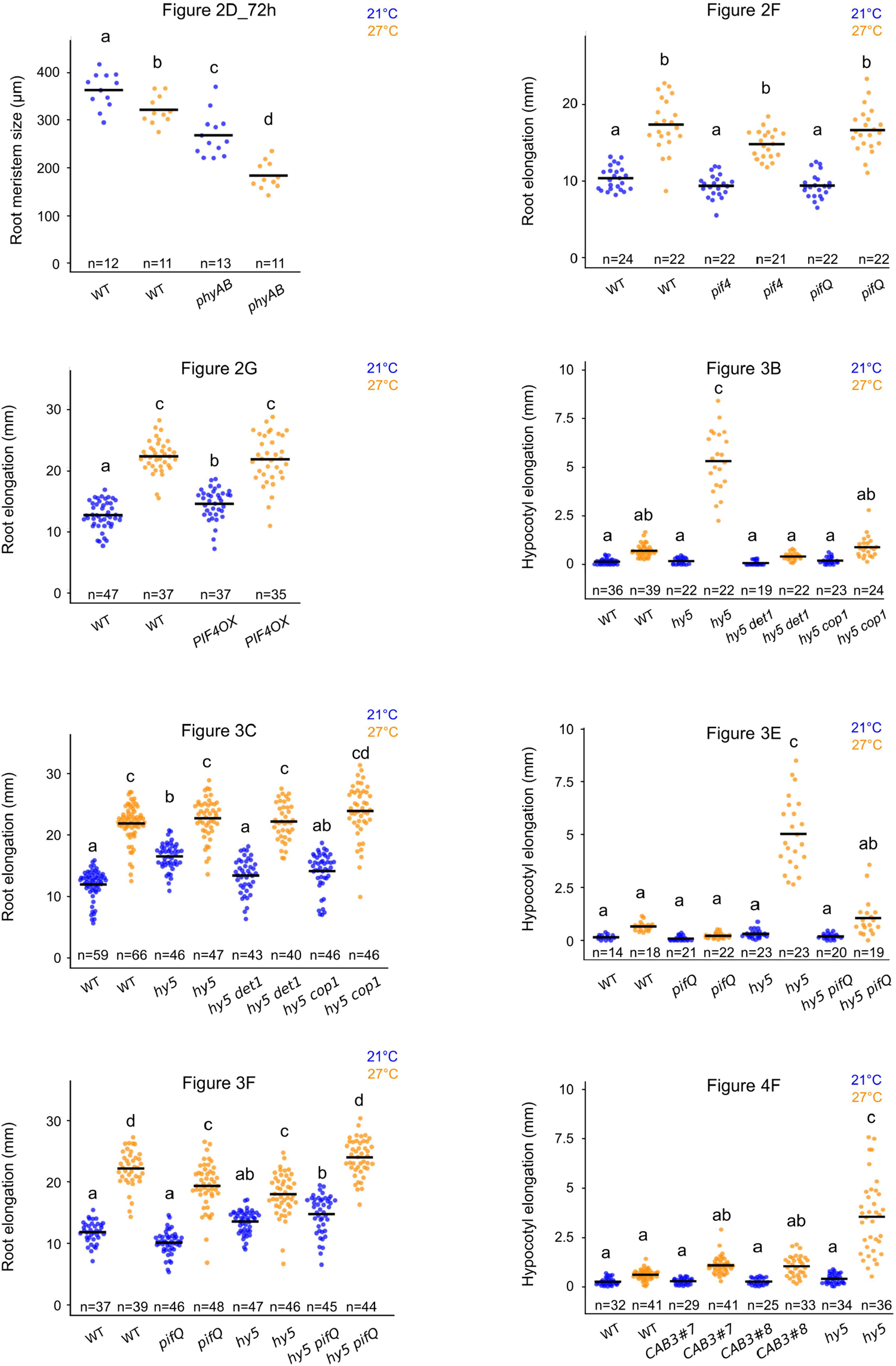

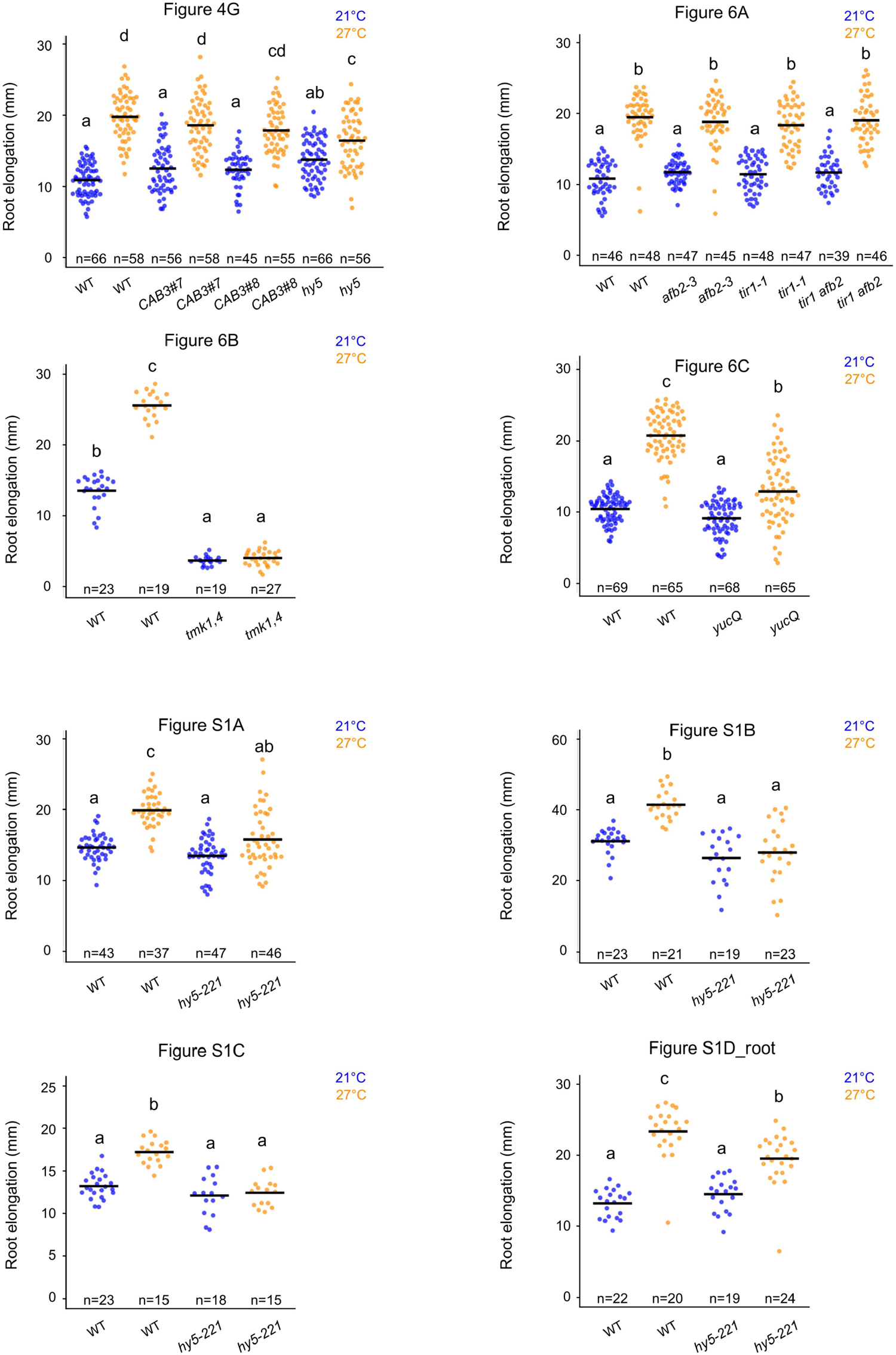

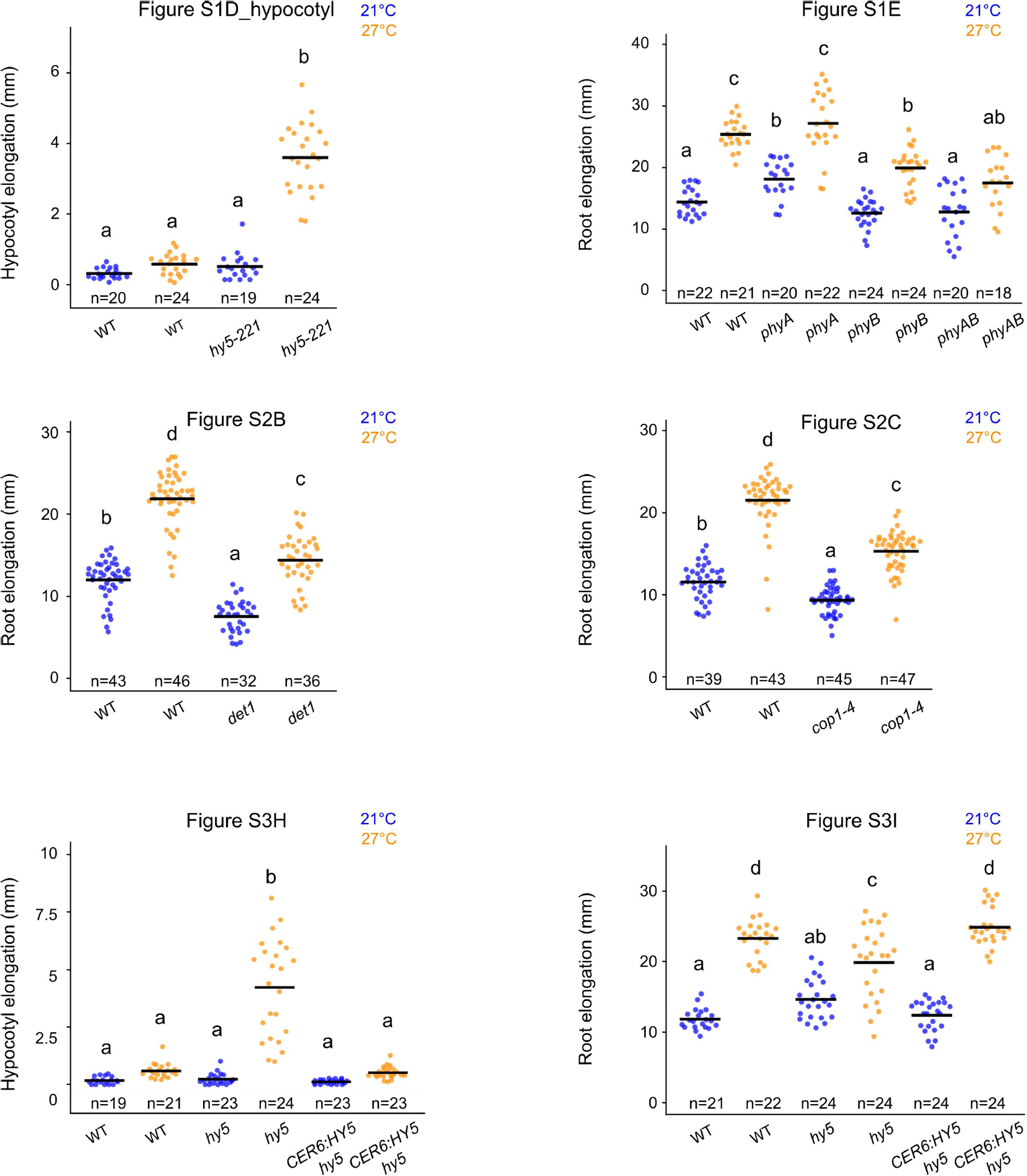

## Notes

### Competing Interest Statement

The authors have declared no competing interest.

### Summary of Updates

Added Source data file

## References

Anders, S., Pyl, P.T., Huber, W., 2014. HTSeq—a Python framework to work with high-throughput sequencing data. Bioinformatics 31, 166–169. https://doi.org/10.1093/bioinformatics/btu638

Bellstaedt, J., Trenner, J., Lippmann, R., Poeschl, Y., Zhang, X., Friml, J., Quint, M., Delker, C., 2019. A Mobile Auxin Signal Connects Temperature Sensing in Cotyledons with Growth Responses in Hypocotyls. Plant Physiol. 180, 757. https://doi.org/10.1104/pp.18.01377

Benjamins, R., Scheres, B., 2008. Auxin: The Looping Star in Plant Development. Annu. Rev. Plant Biol. 59, 443–465. https://doi.org/10.1146/annurev.arplant.58.032806.103805

Burko, Y., Seluzicki, A., Zander, M., Pedmale, U.V., Ecker, J.R., Chory, J., 2020. Chimeric Activators and Repressors Define HY5 Activity and Reveal a Light-Regulated Feedback Mechanism. Plant Cell 32, 967. https://doi.org/10.1105/tpc.19.00772

Burko, Y., Gaillochet, C., Seluzicki, A., Chory, J., Busch, W. 2020b. Local HY5 activity mediates hypocotyl growth and shoot-to-root communication. Biorxiv

Cao, M., Chen, R., Li, P., Yu, Y., Zheng, R., Ge, D., Zheng, W., Wang, X., Gu, Y., Gelová, Z., Friml, J., Zhang, H., Liu, R., He, J., Xu, T., 2019. TMK1-mediated auxin signalling regulates differential growth of the apical hook. Nature 568, 240–243. https://doi.org/10.1038/s41586-019-1069-7

Chen, A., Komives, E.A., Schroeder, J.I., 2006. An Improved Grafting Technique for Mature Arabidopsis Plants Demonstrates Long-Distance Shoot-to-Root Transport of Phytochelatins in Arabidopsis. Plant Physiol. 141, 108. https://doi.org/10.1104/pp.105.072637

Chen, Q., Dai, X., De-Paoli, H., Cheng, Y., Takebayashi, Y., Kasahara, H., Kamiya, Y., Zhao, Y., 2014. Auxin Overproduction in Shoots Cannot Rescue Auxin Deficiencies in Arabidopsis Roots. Plant Cell Physiol. 55, 1072–1079. https://doi.org/10.1093/pcp/pcu039

Chen, X., Yao, Q., Gao, X., Jiang, C., Harberd, N.P., Fu, X., 2016. Shoot-to-Root Mobile Transcription Factor HY5 Coordinates Plant Carbon and Nitrogen Acquisition. Curr. Biol. 26, 640–646. https://doi.org/10.1016/j.cub.2015.12.066

Ciolfi, A., Sessa, G., Sassi, M., Possenti, M., Salvucci, S., Carabelli, M., Morelli, G., Ruberti, I., 2013. Dynamics of the Shade-Avoidance Response in Arabidopsis. Plant Physiol. 163, 331–353. https://doi.org/10.1104/pp.113.221549

Dai, N., Wang, W., Patterson, S.E., Bleecker, A.B., 2013. The TMK Subfamily of Receptor-Like Kinases in Arabidopsis Display an Essential Role in Growth and a Reduced Sensitivity to Auxin. PLOS ONE 8, 1–12. https://doi.org/10.1371/journal.pone.0060990

Delker, C., Sonntag, L., James, G.V., Janitza, P., Ibañez, C., Ziermann, H., Peterson, T., Denk, K., Mull, S., Ziegler, J., Davis, S.J., Schneeberger, K., Quint, M., 2014. The DET1-COP1-HY5 Pathway Constitutes a Multipurpose Signaling Module Regulating Plant Photomorphogenesis and Thermomorphogenesis. Cell Rep. 9, 1983–1989. https://doi.org/10.1016/j.celrep.2014.11.043

Donohue, K., Rubio de Casas, R., Burghardt, L., Kovach, K., Willis, C.G., 2010. Germination, Postgermination Adaptation, and Species Ecological Ranges. Annu. Rev. Ecol. Evol. Syst. 41, 293–319. https://doi.org/10.1146/annurev-ecolsys-102209-144715

Feraru, E., Feraru, M.I., Barbez, E., Waidmann, S., Sun, L., Gaidora, A., Kleine-Vehn, J., 2019. PILS6 is a temperature-sensitive regulator of nuclear auxin input and organ growth in Arabidopsis thaliana. Proc. Natl. Acad. Sci. 116, 3893. https://doi.org/10.1073/pnas.1814015116

Franklin, K.A., Lee, S.H., Patel, D., Kumar, S.V., Spartz, A.K., Gu, C., Ye, S., Yu, P., Breen, G., Cohen, J.D., Wigge, P.A., Gray, W.M., 2011. PHYTOCHROME-INTERACTING FACTOR (PIF4) regulates auxin biosynthesis at high temperature. Proc. Natl. Acad. Sci. 108, 20231–20235. https://doi.org/10.1073/pnas.1110682108

Gangappa, S.N., Botto, J.F., 2016. The Multifaceted Roles of HY5 in Plant Growth and Development. Mol. Plant 9, 1353–1365. https://doi.org/10.1016/j.molp.2016.07.002

Gangappa, S.N., Kumar, S.V., 2017. DET1 and HY5 Control PIF4-Mediated Thermosensory Elongation Growth through Distinct Mechanisms. Cell Rep. 18, 344–351. https://doi.org/10.1016/j.celrep.2016.12.046

Gray, W.M., Östin, A., Sandberg, G., Romano, C.P., Estelle, M., 1998. High temperature promotes auxin-mediated hypocotyl elongation in <em>Arabidopsis</em>. Proc Natl Acad Sci USA 95, 7197. https://doi.org/10.1073/pnas.95.12.7197

Ha, J.-H., Han, S.-H., Lee, H.-J., Park, C.-M., 2017. Environmental Adaptation of the Heterotrophic-to-Autotrophic Transition: The Developmental Plasticity of Seedling Establishment. Crit. Rev. Plant Sci. 36, 128–137. https://doi.org/10.1080/07352689.2017.1355661

Hacham, Y., Holland, N., Butterfield, C., Ubeda-Tomas, S., Bennett, M.J., Chory, J., Savaldi-Goldstein, S., 2011. Brassinosteroid perception in the epidermis controls root meristem size. Development 138, 839. https://doi.org/10.1242/dev.061804

Hanzawa, T., Shibasaki, K., Numata, T., Kawamura, Y., Gaude, T., Rahman, A., 2013. Cellular Auxin Homeostasis under High Temperature Is Regulated through a SORTING NEXIN1-Dependent Endosomal Trafficking Pathway. Plant Cell 25, 3424–3433. https://doi.org/10.1105/tpc.113.115881

Hoecker, U., 2017. The activities of the E3 ubiquitin ligase COP1/SPA, a key repressor in light signaling. Curr. Opin. Plant Biol. 37, 63–69. https://doi.org/10.1016/j.pbi.2017.03.015

Illston, B.G., Fiebrich, C.A., 2017. Horizontal and vertical variability of observed soil temperatures. Geosci. Data J. 4, 40–46. https://doi.org/10.1002/gdj3.47

Jia, K.-P., Luo, Q., He, S.-B., Lu, X.-D., Yang, H.-Q., 2014. Strigolactone-Regulated Hypocotyl Elongation Is Dependent on Cryptochrome and Phytochrome Signaling Pathways in Arabidopsis. Mol. Plant 7, 528–540. https://doi.org/10.1093/mp/sst093

Jung, J.-H., Domijan, M., Klose, C., Biswas, S., Ezer, D., Gao, M., Khattak, A.K., Box, M.S., Charoensawan, V., Cortijo, S., Kumar, M., Grant, A., Locke, J.C.W., Schäfer, E., Jaeger, K.E., Wigge, P.A., 2016. Phytochromes function as thermosensors inseparability. Science 354, 886. https://doi.org/10.1126/science.aaf6005

Kang, Y.H., Breda, A., Hardtke, C.S., 2017. Brassinosteroid signaling directs formative cell divisions and protophloem differentiation in <em>Arabidopsis</em>root meristems. Development 144, 272. https://doi.org/10.1242/dev.145623

Kim, D., Pertea, G., Trapnell, C., Pimentel, H., Kelley, R., Salzberg, S.L., 2013. TopHat2: accurate alignment of transcriptomes in the presence of insertions, deletions and gene fusions. Genome Biol. 14, R36. https://doi.org/10.1186/gb-2013-14-4-r36

Kircher, S., Schopfer, P., 2012. Photosynthetic sucrose acts as cotyledon-derived long-distance signal to control root growth during early seedling development in Arabidopsis. Proc. Natl. Acad. Sci. 109, 11217. https://doi.org/10.1073/pnas.1203746109

Koini, M.A., Alvey, L., Allen, T., Tilley, C.A., Harberd, N.P., Whitelam, G.C., Franklin, K.A., 2009. High Temperature-Mediated Adaptations in Plant Architecture Require the bHLH Transcription Factor PIF4. Curr. Biol. 19, 408–413. https://doi.org/10.1016/j.cub.2009.01.046

Kumar, S.V., Lucyshyn, D., Jaeger, K.E., Alós, E., Alvey, E., Harberd, N.P., Wigge, P.A., 2012. Transcription factor PIF4 controls the thermosensory activation of flowering. Nature 484, 242–245. https://doi.org/10.1038/nature10928

Lau, O.S., Deng, X.W., 2012. The photomorphogenic repressors COP1 and DET1: 20 years later. Trends Plant Sci. 17, 584–593. https://doi.org/10.1016/j.tplants.2012.05.004

Lee, J., He, K., Stolc, V., Lee, H., Figueroa, P., Gao, Y., Tongprasit, W., Zhao, H., Lee, I., Deng, X.W., 2007. Analysis of Transcription Factor HY5 Genomic Binding Sites Revealed Its Hierarchical Role in Light Regulation of Development. Plant Cell 19, 731. https://doi.org/10.1105/tpc.106.047688

Legris, M., Klose, C., Burgie, E.S., Rojas, C.C.R., Neme, M., Hiltbrunner, A., Wigge, P.A., Schäfer, E., Vierstra, R.D., Casal, J.J., 2016. Phytochrome B integrates light and temperature signals in Arabidopsis. Science 354, 897. https://doi.org/10.1126/science.aaf5656

Leivar, P., Monte, E., Oka, Y., Liu, T., Carle, C., Castillon, A., Huq, E., Quail, P.H., 2008. Multiple Phytochrome-Interacting bHLH Transcription Factors Repress Premature Seedling Photomorphogenesis in Darkness. Curr. Biol. 18, 1815–1823. https://doi.org/10.1016/j.cub.2008.10.058

Li, J., Li, G., Gao, S., Martinez, C., He, G., Zhou, Z., Huang, X., Lee, J.-H., Zhang, H., Shen, Y., Wang, H., Deng, X.W., 2010. Arabidopsis Transcription Factor ELONGATED HYPOCOTYL5 Plays a Role in the Feedback Regulation of Phytochrome A Signaling. Plant Cell 22, 3634–3649. https://doi.org/10.1105/tpc.110.075788

Li, L., Ljung, K., Breton, G., Schmitz, R.J., Pruneda-Paz, J., Cowing-Zitron, C., Cole, B.J., Ivans, L.J., Pedmale, U.V., Jung, H.-S., Ecker, J.R., Kay, S.A., Chory, J., 2012. Linking photoreceptor excitation to changes in plant architecture. Genes Dev. 26, 785–790. https://doi.org/10.1101/gad.187849.112

Lian, H.-L., He, S.-B., Zhang, Y.-C., Zhu, D.-M., Zhang, J.-Y., Jia, K.-P., Sun, S.-X., Li, L., Yang, H.-Q., 2011. Blue-light-dependent interaction of cryptochrome 1 with SPA1 defines a dynamic signaling mechanism. Genes Dev. 25, 1023–1028. https://doi.org/10.1101/gad.2025111

Liao, C.-Y., Smet, W., Brunoud, G., Yoshida, S., Vernoux, T., Weijers, D., 2015. Reporters for sensitive and quantitative measurement of auxin response. Nat. Methods 12, 207.

Lorrain, S., Allen, T., Duek, P.D., Whitelam, G.C., Fankhauser, C., 2008. Phytochrome-mediated inhibition of shade avoidance involves degradation of growth-promoting bHLH transcription factors. Plant J. 53, 312–323. https://doi.org/10.1111/j.1365-313X.2007.03341.x

Martins, S., Montiel-Jorda, A., Cayrel, A., Huguet, S., Roux, C.P.-L., Ljung, K., Vert, G., 2017. Brassinosteroid signaling-dependent root responses to prolonged elevated ambient temperature. Nat. Commun. 8, 309. https://doi.org/10.1038/s41467-017-00355-4

McNellis, T.W., von Arnim, A.G., Araki, T., Komeda, Y., Miséra, S., Deng, X.W., 1994. Genetic and molecular analysis of an allelic series of cop1 mutants suggests functional roles for the multiple protein domains. Plant Cell 6, 487. https://doi.org/10.1105/tpc.6.4.487

Novák, O., Hényková, E., Sairanen, I., Kowalczyk, M., Pospíšil, T., Ljung, K., 2012. Tissue-specific profiling of the Arabidopsis thaliana auxin metabolome. Plant J. 72, 523–536. https://doi.org/10.1111/j.1365-313X.2012.05085.x

Omelyanchuk, N.A., Wiebe, D.S., Novikova, D.D., Levitsky, V.G., Klimova, N., Gorelova, V., Weinholdt, C., Vasiliev, G.V., Zemlyanskaya, E.V., Kolchanov, N.A., Kochetov, A.V., Grosse, I., Mironova, V.V., 2017. Auxin regulates functional gene groups in a fold-change-specific manner in Arabidopsis thaliana roots. Sci. Rep. 7, 2489. https://doi.org/10.1038/s41598-017-02476-8

Osterlund, M.T., Hardtke, C.S., Wei, N., Deng, X.W., 2000. Targeted destabilization of HY5 during light-regulated development of Arabidopsis. Nature 405, 462–466. https://doi.org/10.1038/35013076

Oyama, T., Shimura, Y., Okada, K., 1997. The Arabidopsis HY5 gene encodes a bZIP protein that regulates stimulus-induced development of root and hypocotyl. Genes Dev. 11, 2983–2995. https://doi.org/10.1101/gad.11.22.2983

Park, E., Kim, J., Lee, Y., Shin, J., Oh, E., Chung, W.-I., Liu, J.R., Choi, G., 2004. Degradation of Phytochrome Interacting Factor 3 in Phytochrome-Mediated Light Signaling. Plant Cell Physiol. 45, 968–975. https://doi.org/10.1093/pcp/pch125

Park, E., Kim, Y., Choi, G., 2018. Phytochrome B Requires PIF Degradation and Sequestration to Induce Light Responses across a Wide Range of Light Conditions. Plant Cell 30, 1277. https://doi.org/10.1105/tpc.17.00913

Parry, G., Calderon-Villalobos, L.I., Prigge, M., Peret, B., Dharmasiri, S., Itoh, H., Lechner, E., Gray, W.M., Bennett, M., Estelle, M., 2009. Complex regulation of the TIR1/AFB family of auxin receptors. Proc. Natl. Acad. Sci. 106, 22540. https://doi.org/10.1073/pnas.0911967106

Penfield, S., 2008. Temperature perception and signal transduction in plants. New Phytol. 179, 615–628. https://doi.org/10.1111/j.1469-8137.2008.02478.x

Pepper, A., Delaney, T., Washburnt, T., Poole, D., Chory, J., 1994. DET1, a negative regulator of light-mediated development and gene expression in arabidopsis, encodes a novel nuclear-localized protein. Cell 78, 109–116. https://doi.org/10.1016/0092-8674(94)90577-0

Postma, M., Goedhart, J., 2019. PlotsOfData—A web app for visualizing data together with their summaries. PLOS Biol. 17, e3000202. https://doi.org/10.1371/journal.pbio.3000202

Procko, C., Burko, Y., Jaillais, Y., Ljung, K., Long, J.A., Chory, J., 2016. The epidermis coordinates auxin-induced stem growth in response to shade. Genes Dev. 30, 1529–1541. https://doi.org/10.1101/gad.283234.116

Quint, M., Delker, C., Franklin, K.A., Wigge, P.A., Halliday, K.J., van Zanten, M., 2016. Molecular and genetic control of plant thermomorphogenesis. Nat. Plants 2, 15190. https://doi.org/10.1038/nplants.2015.190

Reed, J.W., Nagatani, A., Elich, T.D., Fagan, M., Chory, J., 1994. Phytochrome A and Phytochrome B Have Overlapping but Distinct Functions in Arabidopsis Development. Plant Physiol. 104, 1139. https://doi.org/10.1104/pp.104.4.1139

Reed, J.W., Nagpal, P., Poole, D.S., Furuya, M., Chory, J., 1993. Mutations in the gene for the red/far-red light receptor phytochrome B alter cell elongation and physiological responses throughout Arabidopsis development. Plant Cell 5, 147. https://doi.org/10.1105/tpc.5.2.147

Robinson, M.D., McCarthy, D.J., Smyth, G.K., 2009. edgeR: a Bioconductor package for differential expression analysis of digital gene expression data. Bioinformatics 26, 139–140. https://doi.org/10.1093/bioinformatics/btp616

Rolauffs, S., Fackendahl, P., Sahm, J., Fiene, G., Hoecker, U., 2012. Arabidopsis *COP1* and *SPA* Genes Are Essential for Plant Elongation But Not for Acceleration of Flowering Time in Response to a Low Red Light to Far-Red Light Ratio. Plant Physiol. 160, 2015–2027. https://doi.org/10.1104/pp.112.207233

Saijo, Y., Sullivan, J.A., Wang, H., Yang, J., Shen, Y., Rubio, V., Ma, L., Hoecker, U., Deng, X.W., 2003. The COP1–SPA1 interaction defines a critical step in phytochrome A-mediated regulation of HY5 activity. Genes Dev. 17, 2642–2647. https://doi.org/10.1101/gad.1122903

Salisbury, F.J., Hall, A., Grierson, C.S., Halliday, K.J., 2007. Phytochrome coordinates Arabidopsis shoot and root development. Plant J. 50, 429–438. https://doi.org/10.1111/j.1365-313X.2007.03059.x

Sassi, M., Lu, Y., Zhang, Y., Wang, J., Dhonukshe, P., Blilou, I., Dai, M., Li, J., Gong, X., Jaillais, Y., Yu, X., Traas, J., Ruberti, I., Wang, H., Scheres, B., Vernoux, T., Xu, J., 2012. COP1 mediates the coordination of root and shoot growth by light through modulation of PIN1- and PIN2-dependent auxin transport in Arabidopsis. Development 139, 3402. https://doi.org/10.1242/dev.078212

Shipley, B., Meziane, D., 2002. The balanced-growth hypothesis and the allometry of leaf and root biomass allocation. Funct. Ecol. 16, 326–331. https://doi.org/10.1046/j.1365-2435.2002.00626.x

Slovak, R., Göschl, C., Su, X., Shimotani, K., Shiina, T., Busch, W., 2014. A Scalable Open-Source Pipeline for Large-Scale Root Phenotyping of Arabidopsis. Plant Cell 26, 2390. https://doi.org/10.1105/tpc.114.124032

Sun, J., Qi, L., Li, Y., Chu, J., Li, C., 2012. PIF4–Mediated Activation of YUCCA8 Expression Integrates Temperature into the Auxin Pathway in Regulating Arabidopsis Hypocotyl Growth. PLoS Genet. 8, e1002594. https://doi.org/10.1371/journal.pgen.1002594

Thornley, J.H.M., 1972. A Balanced Quantitative Model for Root: Shoot Ratios in Vegetative Plants. Ann. Bot. 36, 431–441. https://doi.org/10.1093/oxfordjournals.aob.a084602

Tian, T., Liu, Y., Yan, H., You, Q., Yi, X., Du, Z., Xu, W., Su, Z., 2017. agriGO v2.0: a GO analysis toolkit for the agricultural community, 2017 update. Nucleic Acids Res. 45, W122–W129. https://doi.org/10.1093/nar/gkx382

Toledo-Ortiz, G., Johansson, H., Lee, K.P., Bou-Torrent, J., Stewart, K., Steel, G., Rodríguez-Concepción, M., Halliday, K.J., 2014. The HY5-PIF Regulatory Module Coordinates Light and Temperature Control of Photosynthetic Gene Transcription. PLOS Genet. 10, 1–14. https://doi.org/10.1371/journal.pgen.1004416

Van Gelderen, K., Kang, C., Paalman, R., Keuskamp, D., Hayes, S., Pierik, R., 2018. Far-Red Light Detection in the Shoot Regulates Lateral Root Development through the HY5 Transcription Factor. Plant Cell 30, 101. https://doi.org/10.1105/tpc.17.00771

Vijaybhaskar, V., Subbiah, V., Kaur, J., Vijayakumari, P., Siddiqi, I., 2008. Identification of a root-specific glycosyltransferase from Arabidopsis and characterization of its promoter. J. Biosci. 33, 185–193. https://doi.org/10.1007/s12038-008-0036-5

Wang, R., Zhang, Y., Kieffer, M., Yu, H., Kepinski, S., Estelle, M., 2016. HSP90 regulates temperature-dependent seedling growth in Arabidopsis by stabilizing the auxin co-receptor F-box protein TIR1. Nat. Commun. 7, 10269. https://doi.org/10.1038/ncomms10269

Xiong, Y., McCormack, M., Li, L., Hall, Q., Xiang, C., Sheen, J., 2013. Glucose–TOR signalling reprograms the transcriptome and activates meristems. Nature 496, 181.

Xu, T., Dai, N., Chen, J., Nagawa, S., Cao, M., Li, H., Zhou, Z., Chen, X., De Rycke, R., Rakusová, H., Wang, W., Jones, A.M., Friml, J., Patterson, S.E., Bleecker, A.B., Yang, Z., 2014. Cell Surface ABP1-TMK Auxin-Sensing Complex Activates ROP GTPase Signaling. Science 343, 1025. https://doi.org/10.1126/science.1245125

Yanagawa, Y., Sullivan, J.A., Komatsu, S., Gusmaroli, G., Suzuki, G., Yin, J., Ishibashi, T., Saijo, Y., Rubio, V., Kimura, S., Wang, J., Deng, X.W., 2004. Arabidopsis COP10 forms a complex with DDB1 and DET1 in vivo and enhances the activity of ubiquitin conjugating enzymes. Genes Dev. 18, 2172–2181. https://doi.org/10.1101/gad.1229504

Zheng, X., Wu, S., Zhai, H., Zhou, P., Song, M., Su, L., Xi, Y., Li, Z., Cai, Y., Meng, F., Yang, L., Wang, H., Yang, J., 2013. Arabidopsis Phytochrome B Promotes SPA1 Nuclear Accumulation to Repress Photomorphogenesis under Far-Red Light. Plant Cell 25, 115. https://doi.org/10.1105/tpc.112.1070

